# Comparison of Single-Trait and Multi-Trait Genome-Wide Association Models and Inclusion of Correlated Traits in the Dissection of the Genetic Architecture of a Complex Trait in a Breeding Program

**DOI:** 10.1101/2021.08.23.457367

**Authors:** Lance F. Merrick, Adrienne B. Burke, Zhiwu Zhang, Arron H. Carter

**Author notes:** Correspondence: Dr. Arron H. Carter.

## Abstract

Traits with an unknown genetic architecture make it difficult to create a useful bi-parental mapping population to characterize the genetic basis of the trait due to a combination of complex and pleiotropic effects. Seedling emergence of wheat (*Triticum aestivum* L.) from deep planting is a vital factor affecting stand establishment and grain yield, has a poorly understood genetic architecture, and is historically correlated with coleoptile length. This study aimed to dissect the genetic architecture of seedling emergence while accounting for correlated traits using one multitrait GWAS model (MT-GWAS) and three single-trait GWAS models (ST-GWAS). The ST-GWAS models included one single-locus model (MLM), and two multi-locus models (FarmCPU and BLINK). We conducted the GWAS using two populations, the first consisting of 473 varieties from a diverse association mapping panel phenotyped from 2015-2019, and the other population used as a validation population consisting of 279 breeding lines phenotyped in 2015 in Lind, WA, with 40,368 markers. We also compared the inclusion of coleoptile length and markers associated with reduced height as covariates in our ST-GWAS models. ST-GWAS found 107 significant markers across 19 chromosomes, while MT-GWAS found 82 significant markers across 14 chromosomes. FarmCPU and BLINK models were able to identify many small effect markers, and the inclusion of covariates helped to identify large effect markers on chromosome 5A. By using multi-locus models combined with pleiotropic covariates, breeding programs can uncover the complex nature of traits to help identify candidate genes and the underlying architecture of a trait, such as seedling emergence.

## Introduction

Complex traits are controlled by many quantitative trait loci (QTLs) and influenced by environmental conditions (Bernardo 2020). Challenges due to complexity, small effect QTLs, and difficulty in phenotyping, can make it difficult to create useful bi-parental mapping populations to characterize the genetic basis of traits using QTLs. These challenges for linkage mapping for complex traits can result in inconsistent estimated QTL effects (Bernardo 2020; Tibbs Cortes et al. 2021). The difficulty in creating bi-parental mapping populations creates a need for plant breeders and geneticists: to increase the knowledge of the inheritance and genetic architecture of the trait and to identify markers using genome-wide association studies (GWAS; Bernardo 2020).

GWAS enables the discovery of QTLs in a collection of diverse populations or diversity panels rather than using a mapping population (Lander and Schork 1994). GWAS can dissect the genetic architecture of a trait by exploiting all historical recombination events in the population and allow the ability to understand the genetic basis by identifying the associations between genetic markers and phenotypes (Lipka et al. 2015). Not only are complex traits influenced by the environment and multiple QTLs, but they also interact with correlated traits that result in a complex genetic architecture. Using covariates and correlated secondary traits can help account for confounding factors that bias marker effects and improve the GWAS model’s power (Aschard et al. 2015). Commonly used covariates in GWAS account for population structure or genetic relatedness (Tibbs Cortes et al. 2021).

Statistical models have been developed for GWAS to distinguish real associations from false positives caused by population structure and linkage disequilibrium. Some of the first GWAS models, such as general linear models and mixed linear models (MLM), were single-locus, single-trait GWAS (ST-GWAS) models created to implement the covariates along with kinship matrices (Yu et al. 2006). However, these simple models resulted in false negatives caused by weakened associations in order to control inflation of p-values due to population structure (Liu et al. 2016). The MLM was improved upon by compressed mixed linear models (CMLM), which clusters individuals and uses them as random effects rather than individual genotypic effects (Zhang et al. 2010). CMLM was further improved by using pseudo quantitative trait nucleotides (QTNs) to derive kinship instead of all of the genetic markers. The settlement of MLM under progressively exclusive relationship (SUPER) model sorts markers by association and are then divided into bins with the most significant marker designated as pseudo QTNs and then used to derive a reduced kinship matrix (Wang et al. 2014). All of these methods improved computational efficiency over MLM while increasing statistical power. However, the single-locus models tested each marker one at a time, reducing power and increasing computational speed compared to multi-locus models (Liu et al. 2016).

Newer multi-locus models were then developed to test multiple markers simultaneously (Liu et al. 2016). These multi-locus GWAS models, such as the fixed and random model circulating probability unification (FarmCPU) and Bayesian information and linkage-disequilibrium iteratively nested keyway (BLINK) allow the evaluation of big datasets while also reducing false positives and negatives (Huang et al. 2019). FarmCPU was developed in order to control false positives and confounding effects between testing markers and cofactors iteratively. Associated markers are fitted as cofactors to control false positives for testing the rest of the markers in a fixed-effect model, and then a random effect model is used to select the associated markers. BLINK was developed to increase statistical power and efficiency (Huang et al. 2019). BLINK eliminates the FarmCPU assumption that causal genes are evenly distributed across the genome, which improves speed, due to the optimization of bin size and number no longer being required (Huang et al. 2019). However, due to the intricate and pleiotropic nature of quantitative traits, there is no best model for all situations, and it is recommended to use both single locus and multi-locus models to dissect the unique genetic architecture of complex traits (Tibbs Cortes et al. 2021). This is due to differing biological complexities that may be identified by one method and not another, and depends on the genetic architecture and population structure of the trait (Tibbs Cortes et al. 2021).

Additionally, multi-trait GWAS (MT-GWAS) can be used to analyze multiple traits simultaneously. MT-GWAS methods were developed to increase statistical power and identify pleiotropic loci (Porter and O’Reilly 2017). Pleiotropy occurs when genetic loci affect more than one trait (Solovieff et al. 2013). Therefore, correlations and pleiotropy are important, and taking them into account can increase power compared to ST-GWAS (Galesloot et al. 2014). There are various MT-GWAS methods ranging from direct to indirect based models. Direct multivariate approaches consist as a multi-trait mixed model (MTMM), MultiPhen, and bayesian models implemented in MV-SNPTEST and MV-BIMBAM which model the effects of genetic loci directly from the phenotypes (Marchini et al. 2007; O’Reilly et al. 2012; Korte et al. 2012; Stephens 2013; Galesloot et al. 2014). Indirect approaches are based on a reduction of the trait dimensions using canonical correlation analysis in MV-PLINK, or based on principal components using PCHAT (Klei et al. 2008). MT-GWAS methods display an increase in power even when the traits display a negative correlation, when only one of the traits is associated with the loci, or when genetic correlations between the traits are weak (Galesloot et al. 2014). Therefore, MT-GWAS is a valuable approach to understand correlated traits.

Seedling emergence in deep-sown winter wheat exhibits a complex genetic architecture with a significant environmental effect which allows us to compare GWAS methods in order to identify marker-trait associations for selection. Seedling emergence is dependent on deep sowing (sowing depths of 10-15 cm) when precipitation is below 150 to 300 mm annually (Schillinger et al. 1998; Mohan et al. 2013). In deep-sowing practices, fast-emerging cultivars are the most desirable to emerge before precipitation events create soil crusting that can dramatically decrease emergence. Wheat seedlings have decreased emergence when they cannot penetrate the soil surface due to crusting or due to the inability to germinate under dry soil conditions. Seedling emergence and therefore stand establishment is a vital factor affecting grain yield and can reduce grain yields by 35 to 40% (Schillinger et al. 1998).

Previous studies have shown a significant positive relationship between coleoptile length and seedling emergence (Allan et al. 1962; Sunderman 1964; Chastain et al. 1995; Schillinger et al. 1998; Botwright et al. 2001; Schillinger 2011). The reduced height (Rht) genes *Rht-B1b* and *Rht-D1b* are mutant alleles that cause the semi-dwarfing stature of wheat (Vogel et al. 1956; Allan et al. 1962). Dwarfing genes are responsible for short stature and have a pleiotropic effect that includes gibberellin insensitivity, coleoptile length, yield, protein content, and disease resistance (Gale and Youssefian 1983; Allan 1989). Semi-dwarf wheat cultivars have improved resistance to lodging and grain yield but reduced coleoptile length by one-half to three-fourths of the standard varieties at the time of their development (Allan et al. 1961; Allan 1989; Mohan et al. 2013). The reduced coleoptile length was due to decreased gibberellic acid response, which reduced cell size and elongation (Allan et al. 1959). Historically, when crusting was present, or other unfavorable conditions, the shorter coleoptiles of semi-dwarf cultivars resulted in poor stand establishment and yield potential (Rebetzke et al. 1999).

Recently, after 60 years of breeding, emergence in modern varieties was shown to have a reduced correlation between emergence and coleoptile length (Mohan et al. 2013). Coleoptile length only accounted for 28% of the variability for seedling emergence, and some lines with short coleoptiles had the best emergence rating. The remaining variability is attributed to many factors that affect seedling emergence, leading to a complex system resulting in stand establishment. As stated previously, the two main scenarios that affect seedling emergence and subsequently stand establishment include adequate seed-zone water potential for the first leaf to reach the surface and the occurrence of surface soil crust that prevents penetration (Schillinger 2011). Successful seedling emergence is also dependent on the force exerted by the first leaf. The first leaf protrudes through the coleoptile and emerges around 10 to 12 days after planted. During this time, the first leaf can be prone to buckling before it emerges through the soil surface, which can be affected by coleoptile diameter, speed of emergence, and the emergence force and lifting capacity of the first leave along with the previously reported coleoptile length (Arndt 1965; Schillinger et al. 2017; Lutcher et al. 2019). These studies showed that the genetic basis of seedling emergence is a complex trait not solely controlled by genes for any one factor and results in a poorly understood genetic architecture that is dependent on the environment to display variation (Schillinger et al. 2017; Lutcher et al. 2019).

This study presents research to assess the genetic architecture of a complex trait using GWAS. This study’s objectives were to (1) compare ST-GWAS and MT-GWAS models for correlated traits and (2) assess the inclusion of fixed effects to explore the genetic architecture of seedling emergence using different types of populations within and across years to account for differences in genetic variation between populations and environmental variation. This study aided in determining the genetic architecture for a complex trait such as seedling emergence in deep-sown winter wheat.

## Material and Methods

### Phenotypic Data

Seedling emergence notes were taken to select varieties under low annual precipitation (∼150 mm annual precipitation) at Washington State University in Lind, WA, planted using a deep-furrow planting system. Emergence notes were taken on two populations within the breeding program. The diverse association mapping panel (DP) represents a diverse panel of lines not selected exclusively in deep-furrow trials. In contrast, the second population is composed of F_3:5_ breeding lines (BL) and represents a population of closely related lines from a single breeding program composed of pedigrees that have been selected for emergence over previous generations. The two populations were used to compare GWAS models. The DP was used as the primary population for genetic dissection with the BL population used as the validating population. The DP was evaluated in 2015, 2017, 2018, and 2019 in Lind, WA (Table 1). The second population consists of an unreplicated trial of the BL that was evaluated in 2015 (Table 1). In 2016, no data were collected for the DP due to significant soil crusting that was severe enough to impede all lines’ seedling emergence.

**Table 1.** Populations for seedling emergence screened in unreplicated trials under moisture stress in Lind, WA from 2015 to 2019.

| Location | Trial | Year | Individuals |
| --- | --- | --- | --- |
| Lind | DP* | 2015 | 473 |
| Lind | DP | 2017 | 473 |
| Lind | DP | 2018 | 473 |
| Lind | DP | 2019 | 473 |
| Lind | F<sub>3:5</sub> | 2015 | 276 |

\*DP: Quality Association Mapping Diversity Panel

Seedling emergence was visually assessed and recorded as a percentage of the total plot that emerged six weeks after planting for each trial. Table 1 summarizes location, population, year, and the number of genotyped individuals. The emergence issue for each trial was attributed to moisture stress. Coleoptile length was also measured for the DP in 2014 and in two replicates in 2016 under greenhouse conditions. Coleoptile length was recorded to the nearest millimeter according to Murphy et al. (2008).

### Phenotypic Adjustments

Adjusted means from the emergence data collected in the unreplicated trials were adjusted using residuals calculated for the unreplicated lines in individual environments and across environments using the modified augmented complete block design model (ACBD; Federer 1956; Goldman 2019). The adjustments were made following the method implemented in Merrick and Carter (2021), with the full model across environments as follows:

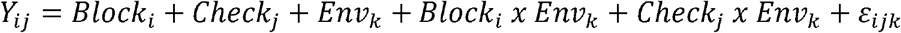

where *Y*_*ij*_ is the trait of interest; *Block*_*i*_ is the fixed effect of the ith block; *Check*_*j*_ is the fixed effect of the jth replicated check cultivar; *Env*_*k*_ is the fixed effect of the kth environment; and ε_*ijk*_ are the residual errors used to adjust the mean of the lines.

Heritability on a genotype-difference basis for broad-sense heritability was calculated using the variance components from the models implemented in Merrick and Carter (2021) and using best linear unbiased predictors for both individual environments and across environments using the formula:

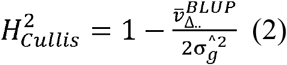

where 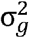, and 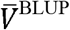 is the genotype variance and mean-variance of the BLUPs (Cullis et al. 2006).

BLUPs for coleoptile length were calculated across trials using a mixed linear model as follows:

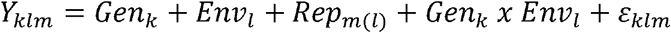

Where *Y*_*klm*_ is the trait of interest; *Gen*_*k*(*j*)_ is the genotype k in the jth Check; *E*_*nvl*_ is the random effect of the lth environment; *Rep*_*m(l)*_ is the replication m in the lth Env; and ε_*ijkl*_ are the residual errors. Trial evaluation and significant differences were evaluated using the coefficient of variation, and analysis of variance (ANOVA) in individual and across trials. Phenotypic correlations were conducted between seedling emergence in the DP across years along with coleoptile length.

Genetic correlations between seedling emergence in the DP and coleoptile length was calculated using the R package “sommer” with the bivariate model, which is represented as:

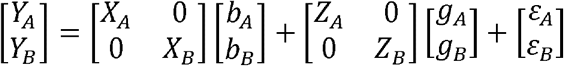

Where *Y*_A_ and *Y*_B_ are adjusted means and BLUPs for emergence and coleoptile length, respectively. X and Z are fixed and random design matrix, subscript A and B represent the primary trait seedling emergence and secondary trait coleoptile length, respectively. a, g, e are the fixed, random genetic effects, and residuals, respectively. Variance components were calculated in the package “sommer”, assuming 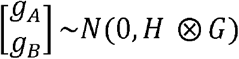 where H is the genetic variance-covariance matrix and G is the additive relationship matrix calculated using “a.mat” function, and 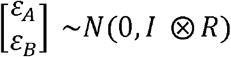 where I is the identity matrix, and R is the residual variance-covariance matrix. Genetic correlations were then calculated using the function “cov2cor” as:

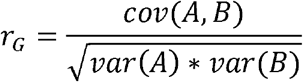

Where cov(A,B) is the covariance between seedling emergence and coleoptile length, and var(A), and var(B) is the genetic variance of seedling emergence and coleoptile length, individually (Covarrubias-Pazaran 2018; R Core Team 2018).

### Genotypic Data

Lines were genotyped using genotyping-by-sequencing (GBS; Elshire et al., 2011) through the North Carolina State Genomics Sciences Laboratory in Raleigh, North Carolina, using the restriction enzymes *Msp*I and *Pst*I (Poland et al. 2012). Genomic DNA was isolated from seedlings at the one to three leaf stage using Qiagen BioSprint 96 Plant kits and the Qiagen BioSprint 96 workstation (Qiagen, Germantown, MD). DNA libraries were prepared following the protocol of DNA digestion with *Pst*I and *Msp*I restriction enzymes (Poland et al, 2012). Genotyping by sequencing (GBS; (Elshire et al. 2011) was conducted at North Carolina State University Genomic Sciences Laboratory with either an Illumina HiSeq 2500 or a NovaSeq 6000. DNA library barcode adapters, DNA library analysis, and sequence SNP calling were provided by the USDA Eastern Regional Small Grains Genotyping Laboratory (Raleigh, NC). Sequences were aligned to the Chinese Spring International Wheat Genome Sequencing Consortium (IWGSC) RefSeq v1.0 (Appels et al. 2018), using the Burrows-Wheeler Aligner (BWA) 0.7.17 (Li and Durbin 2009). Genetic markers with more than 20% missing data, minor allele frequency of less than 5%, and those that were monomorphic were removed. Markers were then imputed using Beagle version 5.0 and filtered once more for markers under a 5% minor allele frequency (Browning et al. 2018). A total of 40,368 single-nucleotide polymorphism (SNP) markers remained.

All winter wheat lines in the DP were genotyped with Kompetitive Allele Specific PCR (KASP®) assays in the WSU Winter Wheat Breeding Laboratory using allele-specific SNP markers for semi-dwarf causing mutant alleles *Rht-B1b* and *Rht-D1b* previously reported in Grogan et al. 2016 and Rasheed et al. 2016. The KASP assays were performed using PACE™ Genotyping Master Mix (3CR Bioscience, Harlow, UK) following the manufacturer’s instructions and endpoint genotyping was conducted from fluorescence using a Lightcycler 480 Instrument II (Roche, Indianpolis, IN). The Rht markers were coded as lines with *Rht-B1b* (1), *Rht-D1b* (2), *Rht-B1b* heterozygous (3), *Rht-D1b* heterozygous (4), *Rht-D1b* with a heterozygous *Rht-B1b* (5), *Rht-B1b* with a heterozygous *Rht-D1b* (6), and both *Rht-B1b* and *Rht-D1b* heterozygous (7).

Linkage disequilibrium (LD) between marker pairs was evaluated using JMP Genomics v.9.0 (SAS Institute, Inc 2011). Significant marker pairs in the same chromosome were considered in LD at a p-value <0.05. Population structure within both populations was analyzed using principal component (PC) analysis biplots and kmeans clustering using the markers in the DP and BL populations individually using the function “prcomp” and “cluster” in R, respectively (R Core Team 2018).

### Genome-Wide Association Models

To dissect the genetic architecture of a complex trait (seedling emergence), ST-GWAS models were implemented using the Genome Association and Prediction Integrated Tool (GAPIT; Liu et al. 2016; Tang et al. 2016; Huang et al. 2019). Both the ST-GWAS and MT-GWAS models were implemented with three principal components fitted as fixed effects. Three principal components were used based on BIC values using model selection in GAPIT (Tang et al. 2016). The GWAS models were conducted on seedling emergence using the adjusted means mentioned previously and on the BLUPs for coleoptile length. The DP was used as the primary population for genetic dissection with the BL population used as the validating population. Three ST-GWAS models were used for comparison. The single-locus ST-GWAS model used was MLM, and the multi-locus models were BLINK and FarmCPU. Within the DP, we compared each model within each year combination without covariates, then with the Rht markers as covariates, coleoptile length BLUPs, and both Rht markers and coleoptile length as covariates. This resulted in 28 datasets for ST-GWAS for seedling emergence in the DP. The GWAS models were then conducted within the BL without covariates to validate the significant markers. In addition, the ST-GWAS models were used to dissect coleoptile length within the DP for further validation to determine whether the significant markers affected coleoptile length.

Additionally, MTMM was implemented using the “sommer” package for MT-GWAS in order to identify pleiotropic interactions between seedling emergence and coleoptile length within the DP. The multivariate Newton-Raphson algorithm was used for multiple random effects and covariance structures using the multivariate model in Covarrubias-Pazaran (2018):

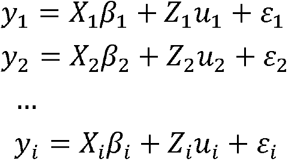

Where y_i_ is a vector of trait phenotypes, β_i_ i is a vector of fixed effects, u_i_ is a vector of random effects for individuals and ε_i_ are residuals for trait “i” (i=1,…,t). The random effects (u_1_ … u_i_ and ε) are assumed to be normally distributed with mean zero. X and Z are incidence matrices for fixed and random effects, respectively. The distribution of the multivariate response and the phenotypic variance covariance V following the models in Covarrubias-Pazaran (2018) are:

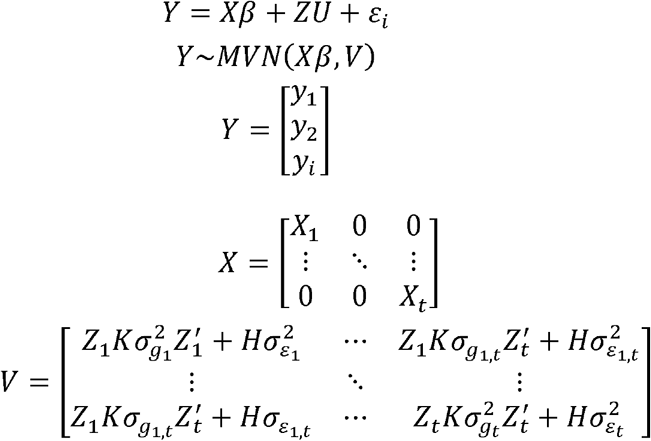

where K is the relationship matrix for the kth random effect (u=1,…,k), and H=I is an identity matrix for the residual term. The terms 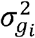 and 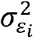 denote the genetic and residual variance of trait “i”, respectively, and σ_*gij*_ and σ_ε*ij*_ are the genetic and residual covariance between traits “i” and “j” (i-1,…,t, and j=1,…,t). Further, the GWAS model implemented in the package “sommer” to obtain marker effects uses the inverse of the phenotypic variance matrix (V) and is a generalized linear model of the form:

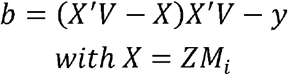

Where: b is the marker effect matrix, y is the multivariate response variable, V-is the inverse of the phenotypic variance matrix, Z is the incidence matrix for the random effect to perform the GWAS, and M_i_ is the ith column of the marker matrix. Further, we implemented three additional F-tests for joint analysis using the scripts proposed in (Korte et al. 2012). The full (FULL) model, which includes the effect of the marker genotype and its interaction, was tested against a null model and identified both loci with common and interaction effects. The interaction model (IE) to identify the interaction effects between the traits tested the full model against a genetic model. Finally, we identified common (COM) genetic effects and tested the genetic model against a null model.

Significant associations based on a Benjamini-Hochberg false discovery rate (FDR; Benjamini and Hochberg 1995). Phenotypic variation and effect explained by each significant marker were calculated by conducting stepwise linear regression between phenotypic and genotypic data and calculating the difference between the effects and variation when a single significant marker was added to the null model in R (R Core Team 2018; Lozada et al. 2019). Significant markers were deemed stable when they were significant in at least two years or in both populations. We then tested the additive effect of pyramiding the stable markers identified across populations and the stable markers identified in multiple years from the DP in each population individually. Manhattan plots were created using the “ggplot2” package, and QQ plots were plotted using the package “CMplots” in R (LiLin 2018; R Core Team 2018).

## Results

### Phenotypic Data

Heritability for seedling emergence was moderately high in the BL and a few years in the DP, while the heritability decreased in combined years. The highest heritability in a single trial was 0.88 in the DP in 2018, and the BL had a heritability of 0.77 (Table 2). The trials with minimum values of zero have a larger standard deviation, which indicates a wider range of seedling emergence and increased environmental pressure and phenotypic variation for selection purposes. Heritability for coleoptile length was very high in the DP (0.89) (Table S1).

**Table 2.**
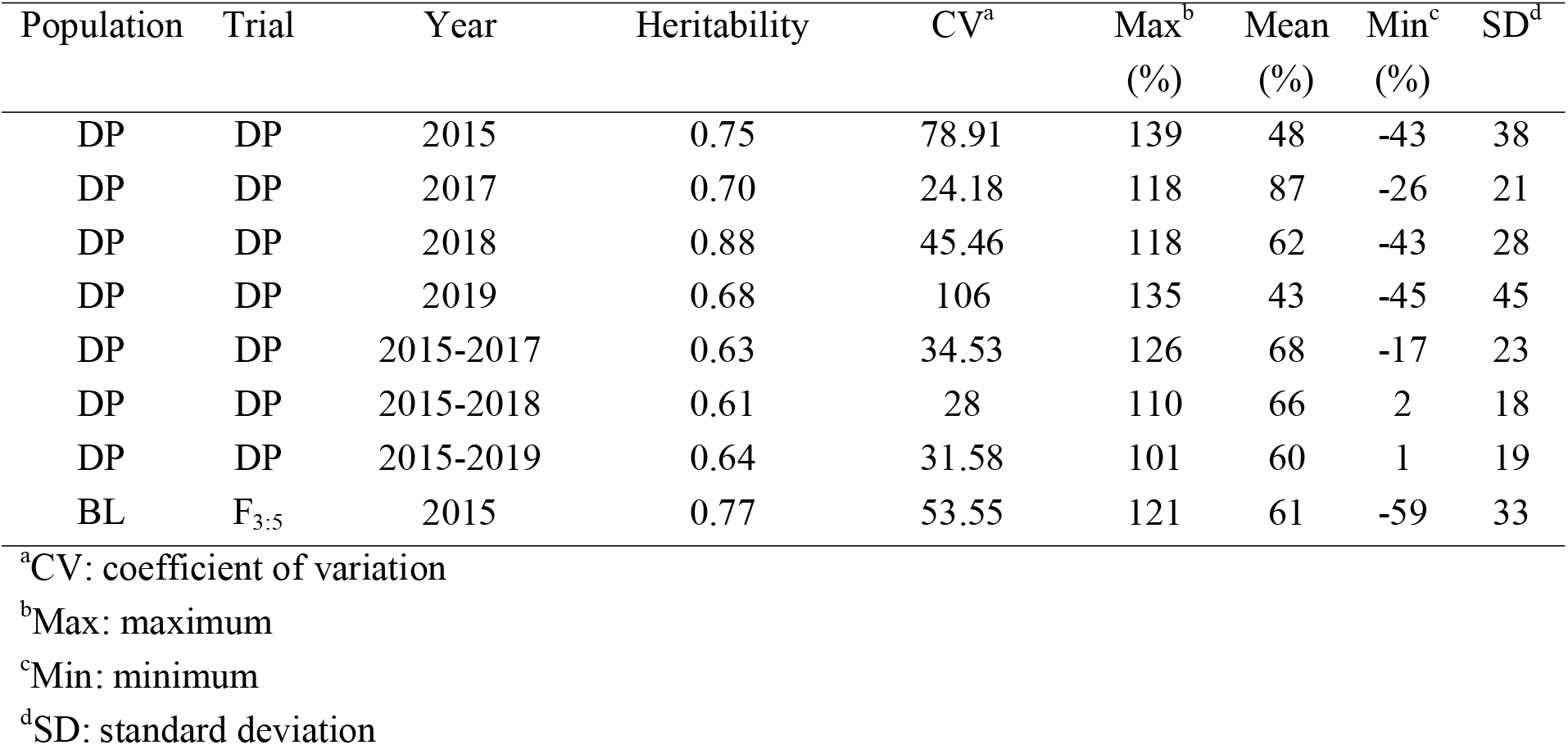
Cullis heritability and trial statistics for deep-sowing seedling emergence in individual and combined trials for the diversity panel (DP) population and breeding line (BL) population phenotyped from 2015 to 2019 and 2015, respectively

Since the variation for seedling emergence depends on the environmental effect, it is important to examine the trials’ variance components. Genetic variances were significant for the DP in 2015, 2017, and for all of the combined trials (Table S2). The environmental effect was not significant in any of the combined trials, with the genotype-by-environment interaction (GEI) effect only significant in the DP over the combined trials of 2015-2018 and 2015-2019 (Table S2). Phenotypic correlations allow us to compare the results in our GWAS models. The DP trials are significantly positively correlated to each other with the exception of three scenarios: DP 2015 to DP 2018; DP 2017 to DP 2018; and DP 2017 to DP 2019 (Table S3). The BL F_3:5_ trial in 2015 was significantly correlated to DP 2017. The genetic correlations between the DP seedling emergence to coleoptile length showed moderate to large correlations in and across years (Table 3). The highest genetic correlation was found in DP 2018 with a correlation of 0.86. However, this was not the case for the phenotypic correlations. The phenotypic correlations in all years was near zero between seedling emergence and coleoptile length (Table 3 and Figure 1). The lack of phenotypic correlation further displays the dependency on the environment for phenotypic variation and the complexity of seedling emergence.

**Table 3.** Genetic correlations between coleoptile length and deep-sowing seedling emergence in individual and combined trials for the diversity panel (DP) population phenotyped from 2015 to 2019

| Year | DP<br>2015 | DP<br>2017 | DP<br>2018 | DP<br>2019 | DP<br>2015-2017 | DP<br>2015-2018 | DP<br>2015-2019 |
| --- | --- | --- | --- | --- | --- | --- | --- |
| Genetic Correlations ($R^2$) | 0.41 | -0.42 | 0.86 | 0.75 | 0.17 | 0.43 | 0.61 |
| Phenotypic Correlations ($R^2$) | 0.00 | -0.01 | -0.06 | 0.03 | 0.00 | -0.04 | -0.01 |

**Fig. 1.**
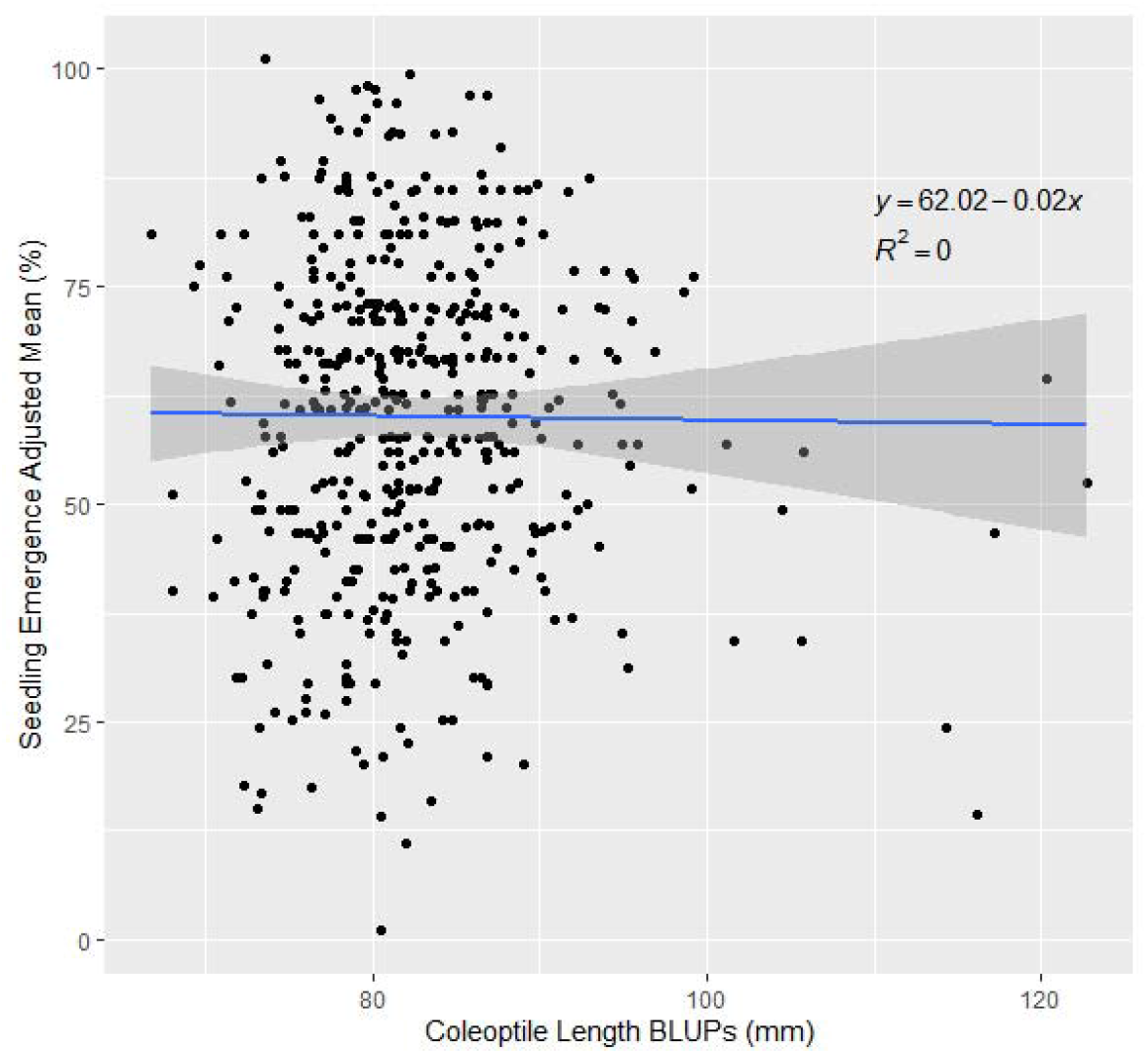
Effect of coleoptile length on seedling emergence in the diversity panel varieties across the combined years of 2015-2019

### Genotypic Data

The principal component biplot using the SNP markers for the DP displayed four clusters according to the elbow method with much overlap between clusters 1 and 2 and a slight overlapping between clusters 3 and 4 (Figure 2A). PC1 explained 12.9% of the variation, and PC2 explained 6.9% of the variation. However, most of the lines clustered within a single cluster in the BL population even though the elbow method revealed four clusters using k-means, and the biplot explained less variation with 9.0% and 5.2% for PC1 and PC2, respectively (Figure 2B). We can visually see the larger genetic variation in the DP than in the BL, and therefore it is important to be accounted for in our GWAS models.

**Fig. 2.**
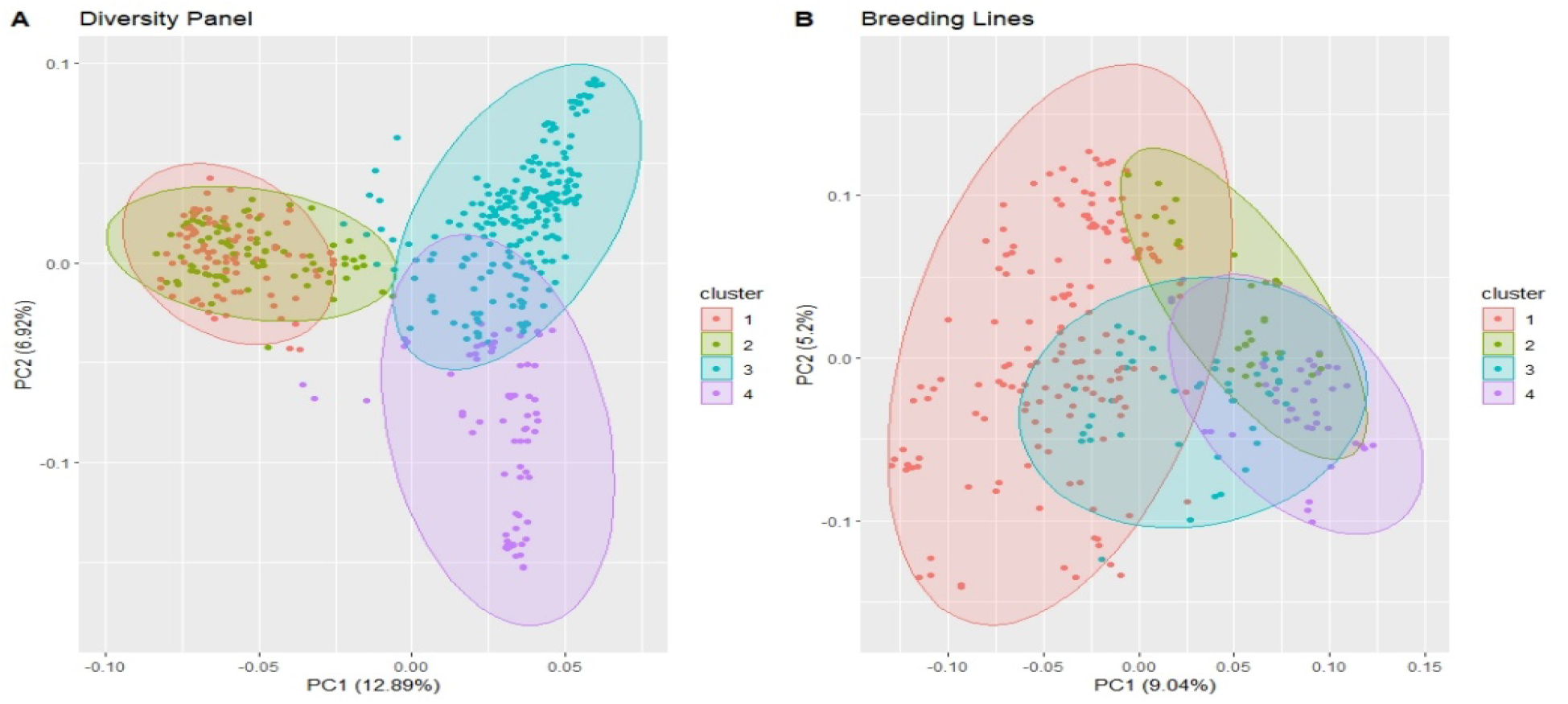
Principal Component (PC) biplot of SNP GBS markers and k-means clustering from the (A) diversity panel and (B) breeding line population consisting of the F_3:5_ trial

The frequency of the Rht alleles in the DP population can be seen in Table S3. The majority of the lines in the DP had either *Rht-B1b (0.564)* or *Rht-D1b* (0.347), with *Rht-B1b* conveying a higher mean seedling emergence than *Rht-D1b* (Figure 3). Further, Figure 3 displays outliers for seedling emergence for lines with certain Rht alleles such as *Rht-D1b*. They were not removed due to the inability to determine the cause of the poor seedling emergence caused by the complexity between phenotypic variation and genotypic effect.

**Fig. 3.**
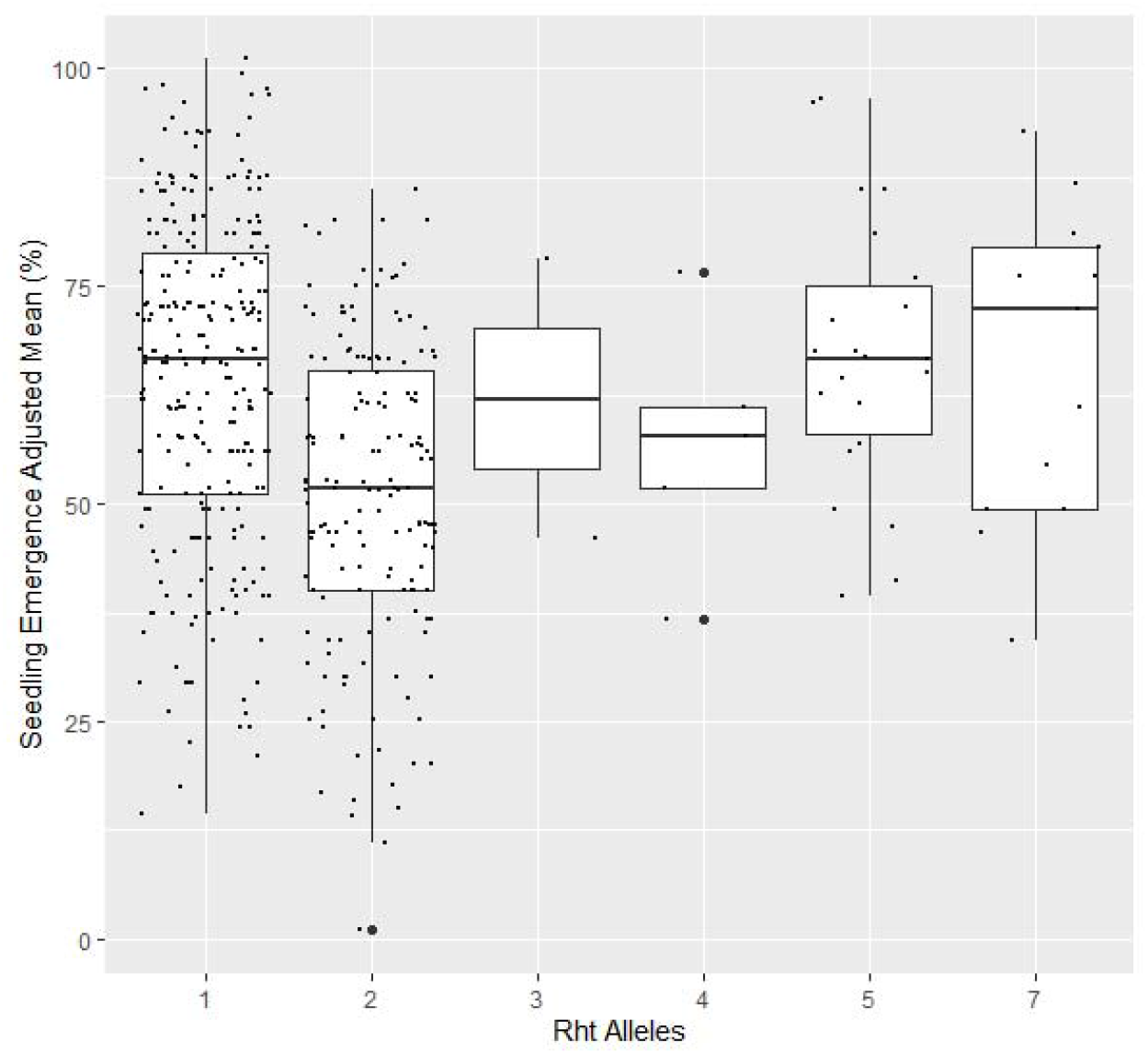
Effect of *Rht* allele frequency on seedling emergence in the diversity panel varieties across the combined years of 2015-2019. The Rht markers were coded as lines with *Rht-B1b* (1), *Rht-D1b* (2), *Rht-B1b* heterozygous (3), *Rht-D1b* heterozygous (4), *Rht-D1b* with a heterozygous *Rht-B1b* (5), *Rht-B1b* with a heterozygous *Rht-D1b* (6), and both *Rht-B1b* and *Rht-D1b* heterozygous (7)

### Single-Trait Genome-Wide Association Studies

To confirm pleiotropic effects of significant markers, ST-GWAS was conducted on coleoptile length within the DP. Genome-wide association for coleoptile length displayed three unique markers using MLM, BLINK, and FarmCPU (Table 4). BLINK and MLM both identified a marker, *S1A_14084576*, on chromosome 1A. Marker *S1A_14084576* conveyed the largest R^2^ value of any significant marker with a value of 7%, and an effect of 14.52. The marker with the next largest R^2^ of 0.05 was *S6A_543395015*. It was identified by both BLINK and FarmCPU on chromosome 6A. It also conveyed the largest effect with 34.44. A third marker was identified only by FarmCPU, *S2A_765677087*, and is located on chromosome 2A. However, *S2A_765677087*, had a rather small effect and R^2^ of 3.22 and 0.04, respectively. The three markers significantly associated with coleoptile length were not identified in any population, year, nor model for seedling emergence (File S1).

**Table 4.** Significant markers for coleoptile length in a Pacific Northwest winter wheat diversity panel using BLINK, FarmCPU and MLM genome-wide association studies (GWAS) models

| Marker | Position<sup>a</sup> | Alleles<sup>b</sup> | Chr<sup>a</sup> | Model | P-value | MAF<sup>c</sup> | Effect (mm)<sup>d</sup> | $R^2$<sup>e</sup> |
| --- | --- | --- | --- | --- | --- | --- | --- | --- |
| *S1A_14084576* | 14084576 | T/A | 1A | BLINK | 1.82E-11 | 0.01 | 14.52 | 0.07 |
|  |  |  |  | MLM | 4.78E-08 | 0.01 | 14.52 | 0.07 |
| S2A_765677087 | 7.66E+08 | G/T | 2A | FarmCPU | 4.22E-09 | 0.14 | 3.23 | 0.04 |
| S6A_543395015 | 5.43E+08 | T/G | 6A | BLINK | 4.70E-08 | 0.00 | 34.44 | 0.05 |
|  |  |  |  | FarmCPU | 6.32E-14 | 0.00 | 34.44 | 0.05 |

<sup>a</sup>Chromosomes and positions of markers were determined according to IWGSC RefSeq v.1.1.
<sup>b</sup>Allele: favorable allele is underlined.
<sup>c</sup>MAF: minor allele frequency.
<sup>d</sup>Effect: Increase of seedling emergence (%) with the inclusion of favorable allele of associated marker.
<sup>e</sup>R<sup>2</sup>: Phenotypic variance explained by associated marker.

There were 107 unique markers over all of the combinations of ST-GWAS for seedling emergence (File S1). Of the 107 significant markers, 96 markers were significant in the DP, and 15 were significant in the BL, with four markers significant in both populations. Seventy-five of the markers were significant in at least two combinations over both populations. Seventy-one and five markers were significant in at least two combinations in the DP and BL, respectively. Significant markers were found on 19 of the 21 chromosomes in the DP, with the majority of the significant markers in the DP located on chromosome 5A with 34 markers (Figure 4). In the BL, only nine chromosomes contained significant markers and more consistent with only 15 total unique markers.

**Fig. 4.**
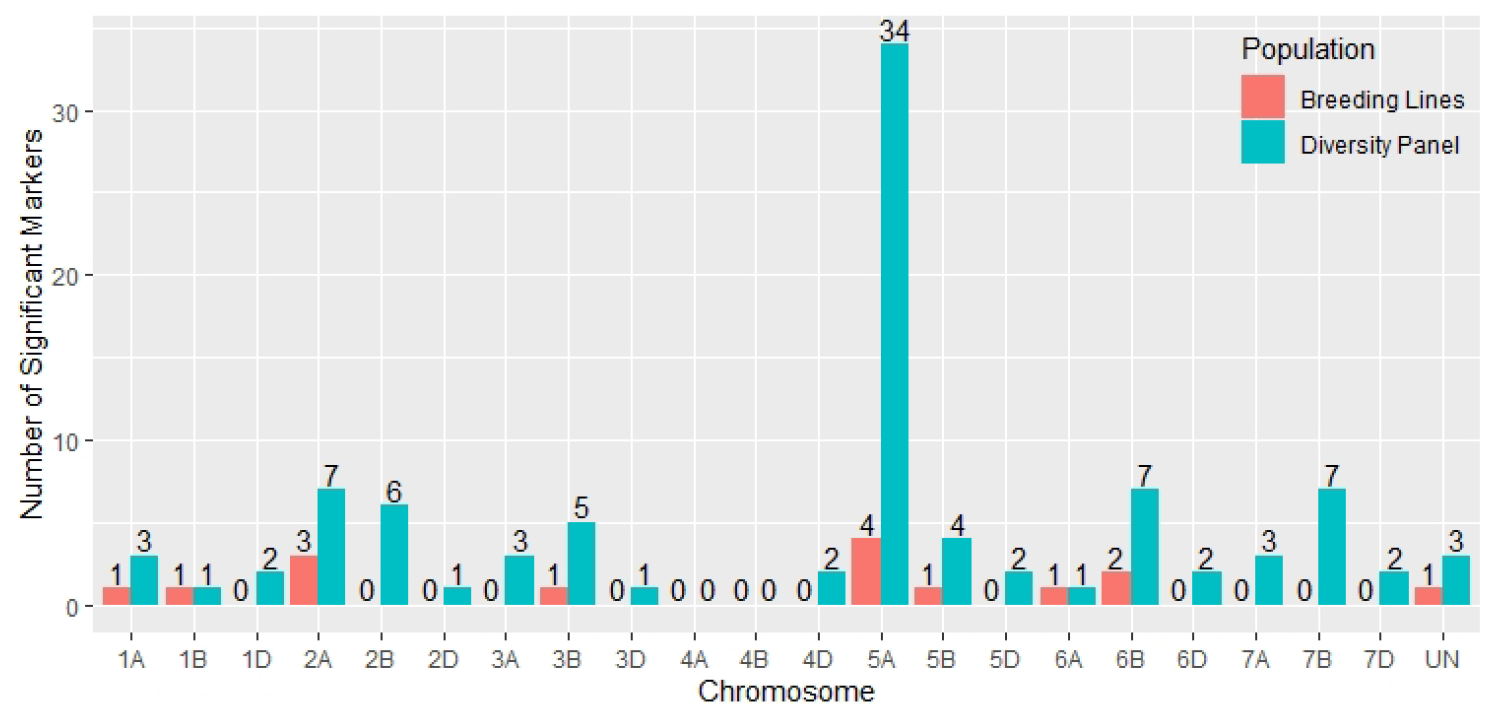
Distribution of significant markers across chromosomes in both the diversity panel varieties and breeding line populations from all combinations of models

All three models had more significant markers in the DP population than in the BL population. The MLM model identified 36 unique markers over all GWAS, three in the BL, and 23 in the DP. FarmCPU identified 65 unique markers overall, nine in the BL, and 57 in the DP. BLINK identified 31 unique significant markers overall, eight in the BL, and 23 in the DP. The MLM model identified significant markers mainly on chromosome 5A with many neighboring significant markers. In contrast, FarmCPU and BLINK identified the same markers on chromosome 5A and identified more markers on various other chromosomes (File S1)

In addition to the effect of the model, there was an indication of the effect in including covariates in BLINK and FarmCPU. This is seen in the number of significant markers identified within the DP (Figure 5). For FarmCPU, including both coleoptile length and Rht alleles individually and combined, decreased the number of significant markers compared to FarmCPU without covariates. For BLINK, the Rht alleles and the combined Rht alleles and coleoptile length increased the number of significant markers, and coleoptile length alone decreased the number of significant markers compared to BLINK without covariates. However, this was not the case for the MLM. For example, including coleoptile length and Rht alleles as covariates alone had similar number of significant markers (64 and 63) compared to the MLM without covariates. Including both Rht alleles and coleoptile length did reduce the number of significant markers to 55. Further, the large number of markers in LD increased the number of significant markers identified with MLM compared to the other ST-GWAS models.

**Fig. 5.**
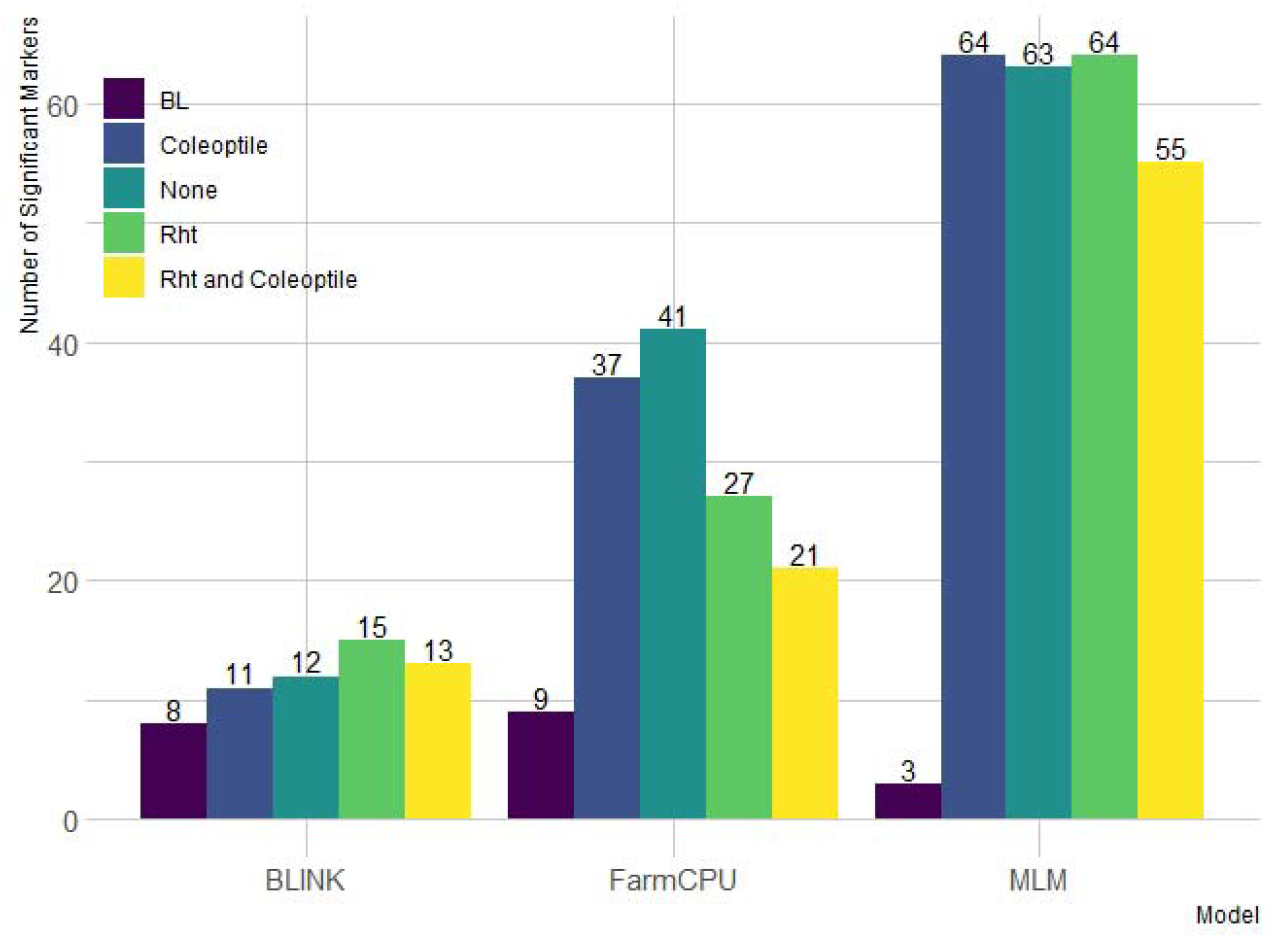
Distribution of significant markers across both the (A) diversity panel and (B) breeding line (BL) populations from all combinations of genome-wide association models. The models compared are BLINK, FarmCPU, and Mixed-Linear Model (MLM) with and without Rht and coleoptile length as covariates in the DP and without covariates in the BL

Further evidence is seen examining QQ plots. For example, in the combined analysis of 2015-2018 and in the BL in 2015, the MLM was consistent regardless of the inclusion of covariates (Figure S1). However, this was not the case with BLINK or FarmCPU. Within the DP, the inclusion of Rht or Rht and coleoptile length increased the deviation from the quantile line more than the models without covariates (Figures S2 and S3). This displays the advantage of using covariates such as Rht alleles within the DP for an increase in power when the trait in question is genetically correlated with another trait. In addition, the QQ plots reveal the difference between the models in the different populations. For both BLINK and FarmCPU, the BL population had the largest deviations compared to the DP GWAS, whereas the opposite was seen in the MLM.

In addition, the environmental impact on the genetic dissection of seedling emergence had a large effect on the identification of significant markers. The GEI for seedling emergence was shown by the varying number of significant markers in different years for the DP. The GWAS models were able to identify the most significant markers in the 2015 trial with 132 markers (Figure S4). However, only a few significant markers were identified in the other individual years with two in 2017 and five in 2019. Further, the combining of years and accounting for GEI in our phenotypic adjustments allowed the GWAS models to increase the ability to dissect seedling emergence compared to individual years. All three year combinations increased the number of significant markers consistently compared to individual years with 109 in year combinations 2015-2017 and 105 in 2015-2018. Additionally, there was a decrease in significant markers identified with the increase in year combinations with the combination of all years (2015-2019) identifying only 47 markers (Figure S4).

### Multi-Trait Genome-Wide Association Studies

The MT-GWAS models were conducted to identify pleiotropic loci between seedling emergence and coleoptile length. The MT-GWAS models displayed very similar results for the individual traits compared to the ST-GWAS models, especially in comparison to the single trait results from the MT-GWAS model. However, we used the Bonferroni cutoff with an alpha =0.05 instead of FDR due to the large deviations and inflation of p-values seen on the QQ plots for the joint models (Figure S5). Using FDR resulted in 924 unique markers across the majority of chromosomes. In comparison, the Bonferroni cutoff resulted in 82 unique significant markers across 14 chromosomes with the majority of the large effect alleles on chromosome 5A (File S2). In comparing the results for MT-GWAS FULL, IE, and COM F-tests to the single trait results from the MT-GWAS, we see no significant markers for coleoptile length significant for any other model (Figure 5). The significant markers for coleoptile length were the same two significant markers on chromosomes 1A and 6A identified in the ST-GWAS models. Additionally, the significant markers for seedling emergence were located on chromosomes 2B (1) and 5A (15).

Further, there were no significant markers in the IE, indicating no contradictory effect markers between seedling emergence and coleoptile length. The lack of significant IE markers is why the FULL model and the COM models are very similar. The COM model was also able to identify significant markers in every year in the DP compared to the single trait results for seedling emergence, which did not identify significant markers in the 2017 through 2019 individual years (File S2). Additionally, there were 64 significant markers in the COM model, with 16 significant markers also found significant for seedling emergence (Figure 6). Therefore, indicating the potential for pleiotropic markers and the ability to identify them using a joint analysis model compared to the single trait results implemented in the MT-GWAS models.

**Fig. 6.**
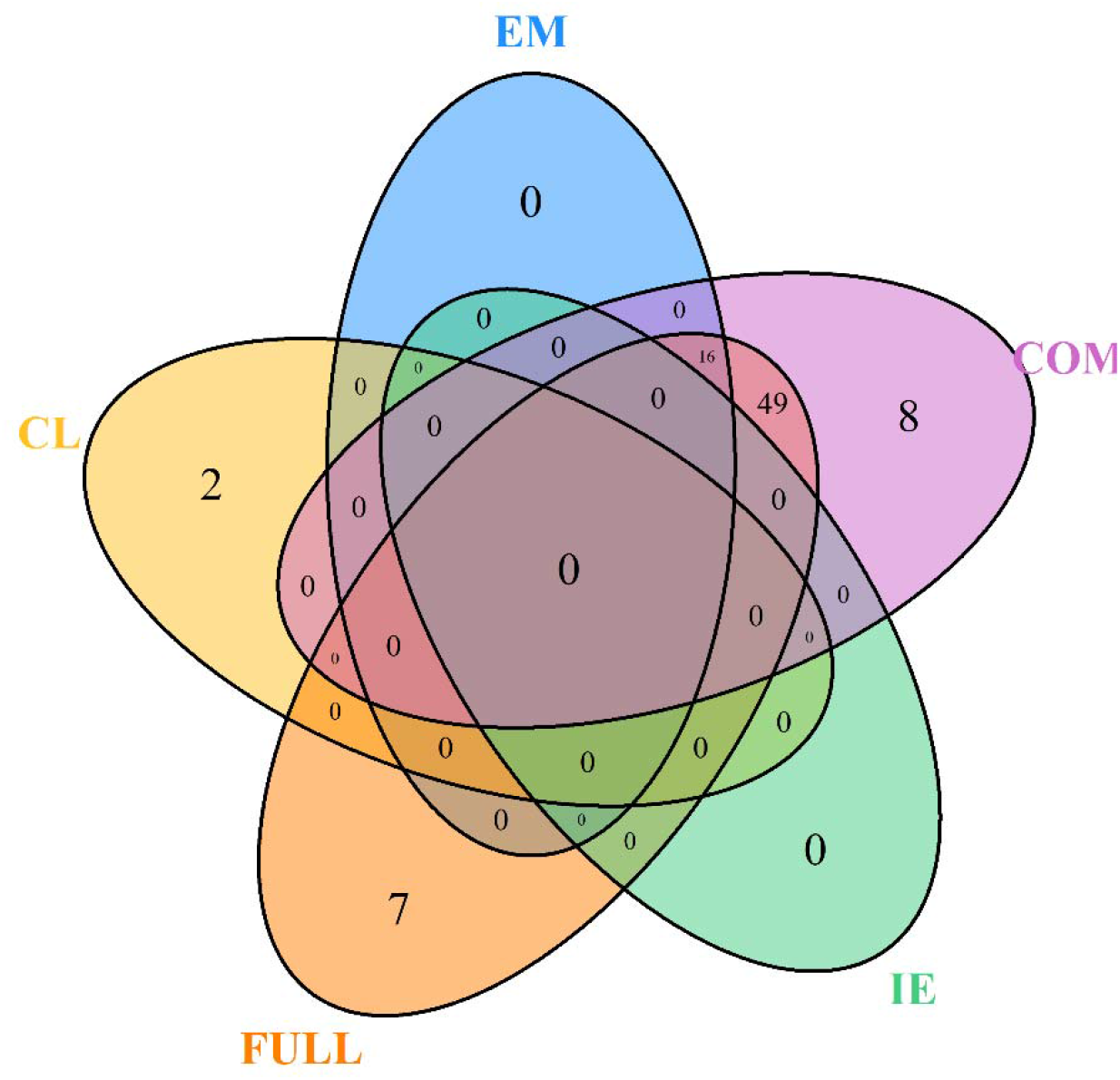
Venn diagram for the number of unique significant markers across the multi-trait genome-wide association studies (MT-GWAS) for seedling emergence (EM), coleoptile length (CL), and the joint analysis for seedling emergence and coleoptile using the full effect (FULL), interaction effect (IE), and common effect (COM) multi-trait mixed models for identifying significant loci controlling deep-sowing seedling emergence and coleoptile length in a Pacific Northwest winter wheat diversity panel (DP) phenotyped across trials from 2015-2019 in Lind, WA

### Stable Significant Markers

To validate the significant markers, it is important to compare their significance across years, populations, and in correlated traits to determine their stability as significantly associated with seedling emergence. As mentioned previously, no markers were found significant in both seedling emergence and coleoptile length in either ST-GWAS or MT-GWAS. In the ST-GWAS models, only 74 of the 107 markers were found significant in more than one GWAS. Within the DP, 23 unique markers were found significant in more than one year across models. BLINK only identified one marker, *S5A_522153944*, in multiple years. FarmCPU identified six markers across years on chromosomes 2A (2), 2B (1), and 5A (3). MLM identified 16 markers across years, with all but one on chromosome 5A with the other marker located on chromosome 5B. For these stable markers in the DP, only *S5A_522153944* was identified by all three models.

Out of the 107 unique significant markers, only four markers were identified in both populations. Further, only *S5A_522153944, S5A_522153953*, and *S5A_523025549* were identified across both populations with the same model (Table 5). MLM identified all three of these markers in both populations. whereas BLINK did not identify any markers in both populations, and FarmCPU only identified *S5A_522153944* in both populations. The effect of covariates on identifying the stable markers was inconsistent in the DP. MLM displayed no response to the inclusion of covariates (Figure S6). However, FarmCPU identified the stable marker *S5A_522153944* on chromosome 5A using the Rht markers, coleoptile length, and Rht markers with coleoptile length combined as covariates (Figure S7). Further, BLINK was able to detect *S5A_522153944* with and without covariates but was not able to identify it in the BL population (Figure S8).

**Table 5.**
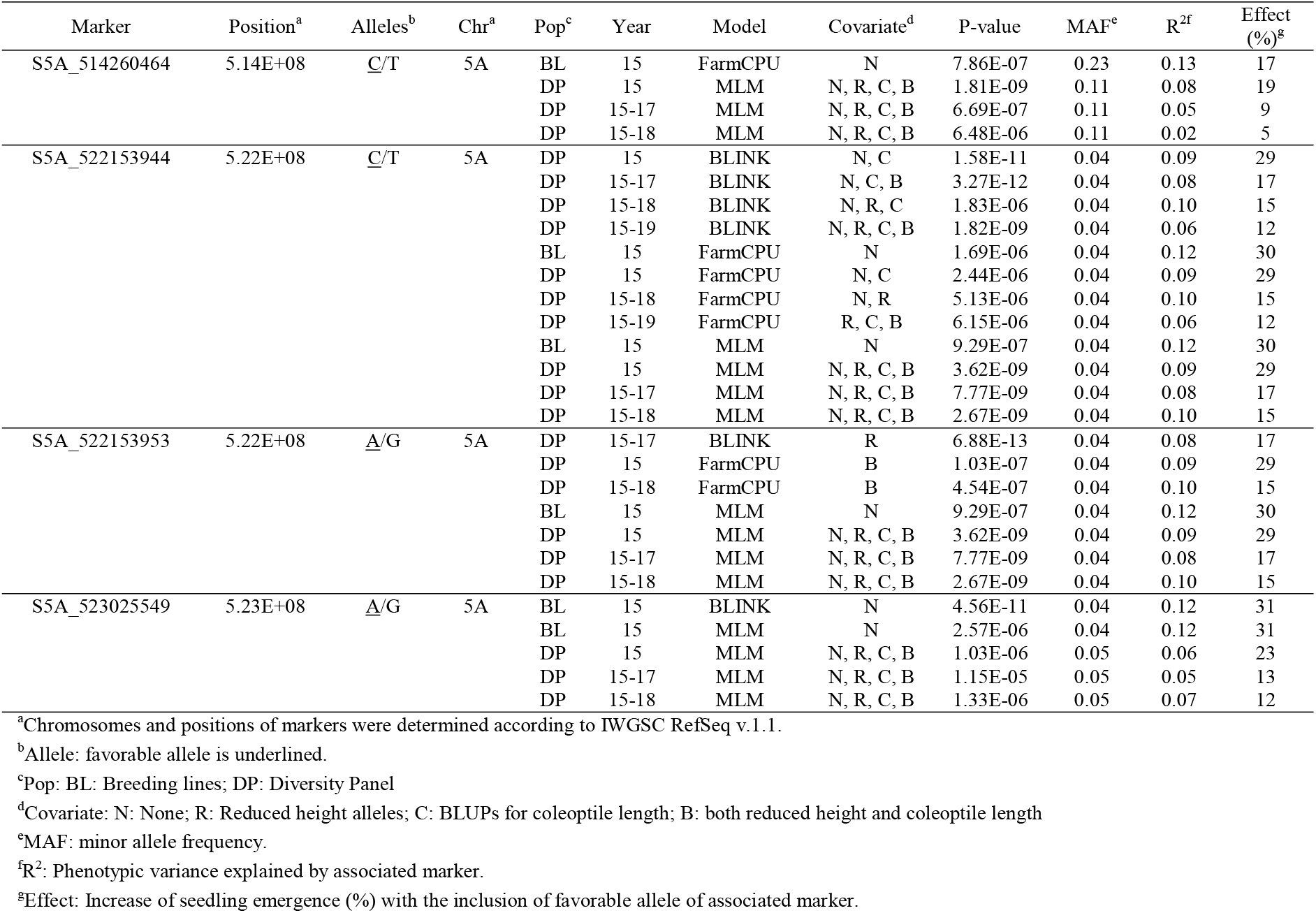
Stable significant markers across both the diversity panel and breeding lines for controlling deep-sowing seedling emergence in a Pacific Northwest winter wheat breeding trial using BLINK, FarmCPU, and MLM genome-wide association studies (GWAS) models using covariates for coleoptile length and Rht alleles

When taking into account the number of significant markers across chromosomes, QQ plots, covariate effect, and across population identification, FarmCPU displayed the ability to identify both the stable large effect and small effect markers with fewer false positives on chromosome 5A (Figure 7). In comparison, MLM consistently identified markers regardless of covariates and population but also identified a large number of linked markers. For example, MLM consistently identified both *S5A_522153944* and *S5A_522153953* which have an R^2^ LD value of 1, and identified 11 other markers on 5A with R^2^ LD values above 0.80. Further, BLINK was able to identify other small effect loci on other chromosomes but did not identify a significant marker in both populations.

**Fig. 7.**
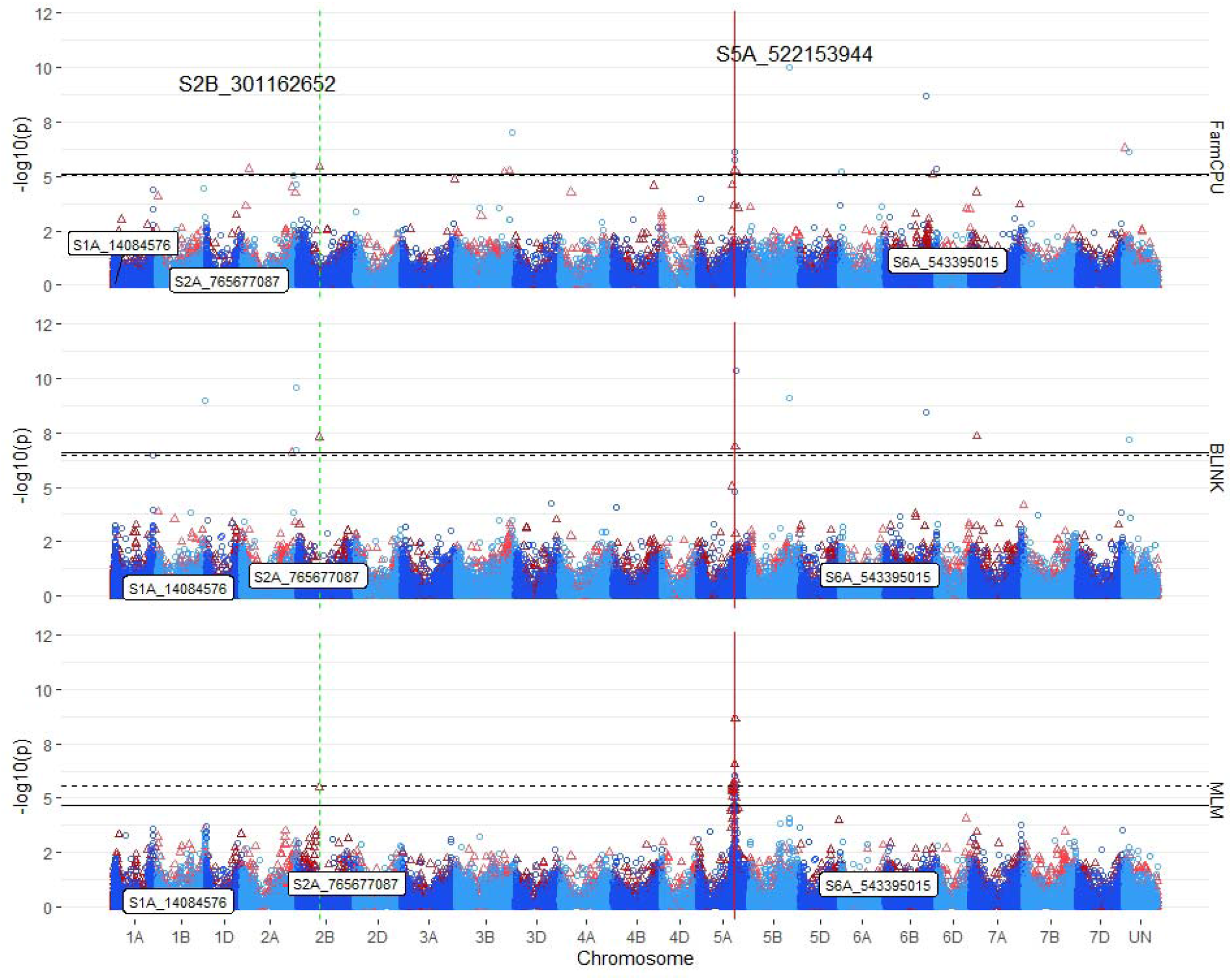
Stacked manhattan plots for genome-wide association studies (GWAS) using MLM, FarmCPU, and BLINK for identifying significant loci controlling deep-sowing seedling emergence in a Pacific Northwest winter wheat diversity panel (DP) phenotyped across trials from 2015-2018 and a breeding line (BL) trial phenotyped in 2015 in Lind, WA. Red triangles display GWAS results in the DP, and blue circles identify GWAS results in the BL. Significant markers using an FDR cutoff with an alpha=0.05 are placed above the solid black line for the DP and the dashed black line for the BL. Significant markers across both populations are identified with a vertical red solid line identifying their positions. Markers enclosed in a white text box display significant markers identified in GWAS studies for coleoptile length

The stable markers identified across years in the DP were located on chromosome 2A (2), 2B (3), 5A (15), 5B (1), and 7A (1). The two largest effect stable markers in the DP, *S5A_522153944* and *S5A_522153953*, were found on chromosome 5A and had the largest effect of 30% for seedling emergence. These two markers are in complete linkage with an R^2^ LD value of 1. They are both considered relatively large effect alleles and had a maximum R^2^ of 10% in the DP. The other stable markers across years identified on chromosome 5A were mainly identified by MLM and their effects ranged from 3% to 29%, with R^2^ values ranging from to 2% to 8%. FarmCPU was able to identify stable markers on the other chromosomes mentioned previously. However, they were minor effect markers with effects ranging from 4 to 24% on seedling emergence and R^2^ values from 1 to 6%. MLM was only able to identify the smaller effect markers on chromosomes 2B and 5B. The multi-locus models, FarmCPU and BLINK, were able to identify the remaining small effect markers primarily in the combined year trials. Overall, the stable markers across years accounted for 30% of the total variation in the combined 2015-2019 DP.

GWAS in the BL population displayed more consistent results than in the DP, and only identified stable markers on chromosome 5A. There were four significant stable markers identified in the BL (Table 5). The four stable markers were all found in the DP, and had effect ranges of 17 to 30%, with R^2^ values of 12 to 13%. The stable markers in the BL population accounted for 18% of the total variation for seedling emergence, displaying a more consistent dissection of the complex trait compared to the DP.

The MT-GWAS models displayed very similar results for identifying stable markers across years for both the single trait and joint models as the ST-GWAS models. The stable markers for seedling emergence and coleoptile length as the ST-GWAS models, with the seedling emergence stable markers being located exclusively on chromosomes 2B and 5A, whereas the stable markers for coleoptile length were the same markers identified in the ST-GWAS models on chromosomes 1A and 6A. In addition to the stable singe trait markers, the COM MT-GWAS model identified 19 stable markers on chromosome 5A, including the large effect marker *S5A_522153944* (Figure 8). In addition, the COM model identified stable markers on chromosomes 2B (1), 5B (1), and 7B (3). The 2B and 5B markers were *S2B_301162652* and *S5B_491273019.* The 7B markers are all completely linked with R^2^ values of 1, with the marker *S7B_663828309* having the largest effect. The 7B markers were the only stable markers in MT-GWAS not found stable in the ST-GWAS models. The 7B markers were only found significant in the 2015-2017 combined DP for the ST-GWAS models, and therefore, were not deemed stable.

**Fig. 8.**
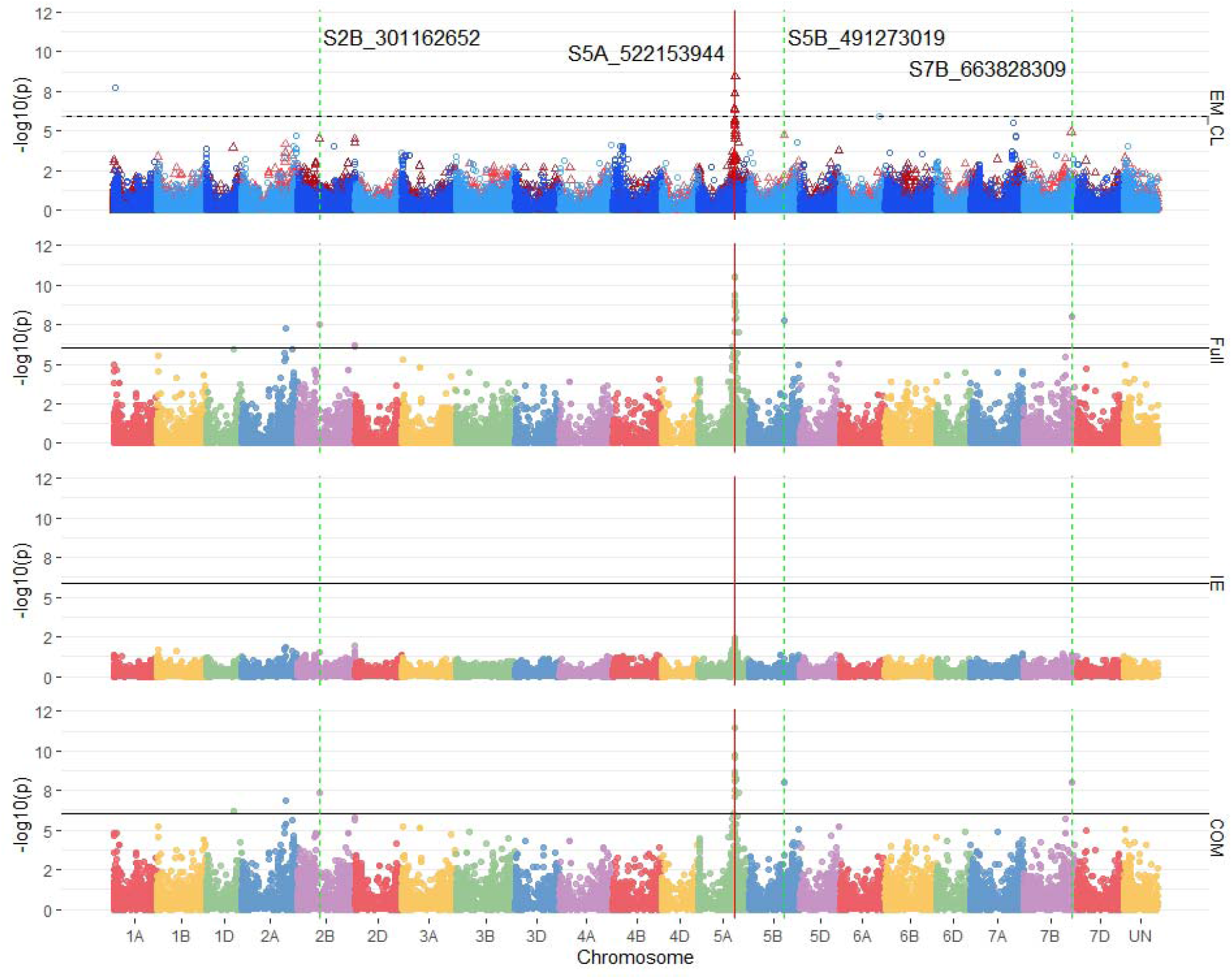
Stacked manhattan plots for multi-trait genome-wide association studies (MT-GWAS) using seedling and emergence overlapped (EM_CL), the full effect (Full), interaction effect (IE), and common effect (COM) multi-trait mixed models for identifying significant loci controlling deep-sowing seedling emergence and coleoptile length in a Pacific Northwest winter wheat diversity panel (DP) phenotyped across trials from 2015-2018 in Lind, WA. For the overlapped plot, EM_CL, red triangles display GWAS results for seedling emergence, and blue circles identify GWAS results for coleoptile length. Significant markers using a Bonferonni cutoff with an alpha=0.05 are placed above the solid black line for seedling emergence and the dashed black line for coleoptile length. Significant markers across three models are highlighted with a solid red vertical line, and significant markers across two models are highlighted with a dashed green vertical line identifying their positions

### Effect of Combining Favorable Alleles

The frequencies of the stable markers across populations and the stable markers across years for seedling emergence are important for understanding the populations’ selection and makeup. The stable markers with LD above 0.80 R^2^ were binned together with the marker with the largest effect identified to represent the bin. This resulted in three out of the four markers identified across populations remaining and 12 out of 23 markers identified across years remaining for displaying their additive effect for seedling emergence (Table S5). In addition, the significant marker on 7B identified in the COM MT-GWAS model was included with the markers stable across years. The majority of stable markers had high frequencies in the 90% range in both the DP and BL (Table S5). This shows that the stable markers are already consistent in both populations.

Even though the stable markers across populations and years were frequent in the populations, the differences was shown when we combined them and compared the favorable alleles’ cumulative effect on seedling emergence (Figure 9A-D). We compared the additive ability for favorable alleles using stable markers across populations and stable markers identified in the DP in multiple years. For the stable markers across populations, the majority of lines in both populations had favorable alleles for all three of the markers, which were all located on chromosome 5A. Both populations showed an additive effect, but in the DP, the lines with all three markers had a lower mean compared to lines with just two of the markers (Figure 9B). However, in the BL there was an increasing trend in emergence with the accumulation in favorable alleles (Figure 9A). The stable markers across the populations only accounted for 8% of the total variation in the DP and 19% in the BL.

**Fig. 9.**
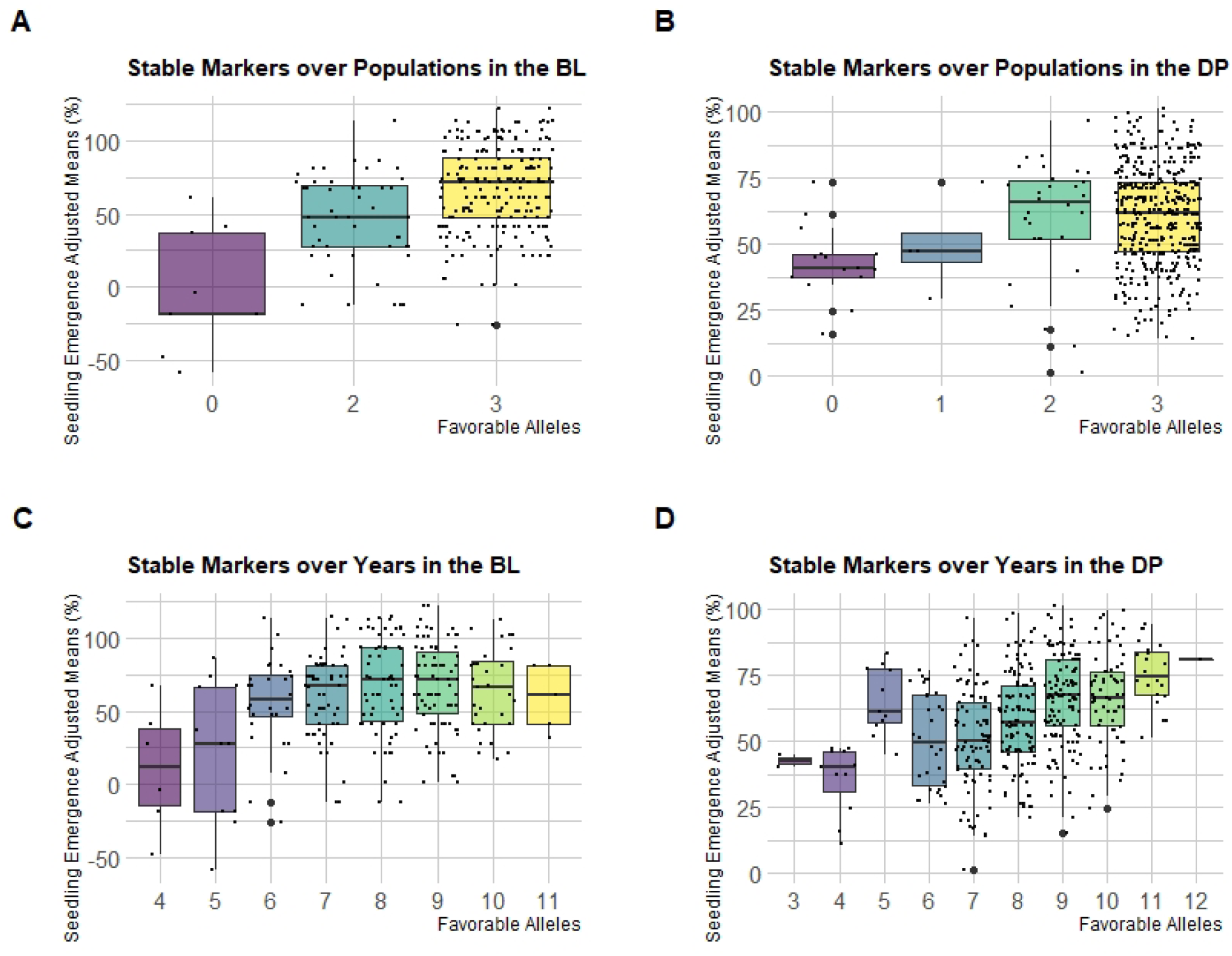
Effect of pyramiding favorable alleles for seedling emergence for the stable markers identified in both populations (A and B), and stable markers identified across years in the DP (C and D) across both the (A and C) diversity panel (DP) and (B and D) breeding line (BL) populations

This additive effect is seen to a point for the stable markers across years for seedling emergence in the BL. The DP showed an additive effect for all 11 favorable alleles, whereas the BL showed a diminishing return signifying some of the markers may not have a large effect within the BL population (Figure 9C and 9D). For the DP, the majority of lines had either eight or nine favorable alleles with 113 and 125 lines, respectively. This was also the case in the BL population with 72 lines having eight alleles and 72 lines having 9 alleles. The stable markers across years accounted for only 23% of the variation in the BL and 31% in the DP.

## Discussion

### Complex Traits and Seedling Emergence

Complex traits are quantitative in nature and are affected by many small-effect QTLs (Holland 2007). The challenges that impede the understanding of complex traits are the inability to statistically detect and map minor effect QTLs, accurately understand GEI, and account for pleiotropic effects (Luo et al. 2017). This study attempted to characterize one such trait, seedling emergence for deep-sown winter wheat, to be used as a model for other complex traits.

The reduced height semi-dwarf genes *Rht-B1b* and *Rht-D1b* are mutant alleles that cause the dwarfing stature of wheat due to the decrease in response to endogenous gibberellic acid (Vogel et al. 1956; Allan et al. 1962; Peng et al. 1999). These dwarfing genes are responsible for short stature and have a pleiotropic effect that includes gibberellin insensitivity, coleoptile length, grain yield, protein content, and disease resistance (Allan et al. 1961; Allan 1989; Mohan et al. 2013). The introgression of these genes into ‘Norin 10’ and the subsequent use by N.E. Borlaug was an essential driver of the green revolution (Vogel et al. 1956; Allan 1989; Mohan et al. 2013). Semi-dwarf wheat cultivars were resistant to lodging, which allowed more fertilizer input and much-improved grain yield (Allan 1989; Mohan et al. 2013). A side effect of the semi-dwarf genes’ introgression was the reduction in coleoptile length of one-half to three-fourths of the standard varieties at the time (Allan et al. 1961). The reduced coleoptile length is due to a decreased response of GA, which reduced cell size and elongation (Allan et al. 1959). Historically, when crusting was present, or other unfavorable conditions, the shorter coleoptiles of semi-dwarf cultivars resulted in poor stand establishment and yield potential (Rebetzke et al. 1999). These previous studies showed the relationship between coleoptile length and seedling emergence (Allan et al. 1962; Sunderman 1964; Chastain et al. 1995; Schillinger et al. 1998; Botwright et al. 2001; Schillinger 2011). However, the study by Mohan et al. (2013) was able to refute the previous claims in modern varieties using a collection of 662 wheat cultivars from around the world. The previous studies had claimed that coleoptile length explained more than 60% of the variation for seedling emergence but only explained 28% in modern varieties (Mohan et al. 2013).

Major QTLs for coleoptile length have been reported on chromosomes 4BS, 4BL, and 5AL in wheat (Rebetzke et al. 2001). The QTLs on 4B were reported on either side of the *Rht-B1* allele. This study was further enhanced by a subsequent QTL analysis that resolved the two 4B QTLs directly to the *Rht-B1* locus (Rebetzke et al. 2007). In our study, no markers were located on 4B with two small effect markers on 4D found only in the joint COM and FULL MT-GWAS models, but were not found stable across years. The major stable markers validated across populations for seedling emergence in our study were identified on chromosome 5A. However, in the follow-up study in Rebetzke et al. (2007), the QTL for coleoptile length on 5A was not found repeatable across two or more populations. There have been small-effect QTLs for coleoptile length identified on wheat chromosomes 1A, 2B, 2D, 3A, 3B, 5A, 6A (Rebetzke et al. 2007). We found significant markers in our GWAS models on these chromosomes, with our large effect markers also located on chromosome 5A. However, in the DP, significant markers for coleoptile length were located on chromosomes 1A, 2A, and 6B, and none of these markers were significant for seedling emergence. Therefore, we can conclude that our GWAS models are not selecting major markers for coleoptile length. Further, we can conclude that the seedling emergence we are dissecting is indeed due to other factors affecting seedling emergence or another poorly understood mechanism similar to what was indicated in Mohan et al. (2013).

Our study demonstrated that chromosome 5A has a large effect on seedling emergence with four large effect stable markers identified consistently across both years in the DP and validated in the BL. Therefore, a major effect QTL for seedling emergence may be present in that chromosomal location. In our GWAS models, the significant markers are possibly affecting both fast emergence and the ability to deal with moisture stress. The variation for emergence in the DP and BL was due to moisture stress, and therefore, the significant markers would affect both fast-emergence and the ability to deal with moisture stress. However, as mentioned previously, seedling emergence is a complex system of multiple factors and pleiotropic effects that are not controlled by genes for any sole component.

The lack of consistency between GWAS models and populations in individual years in the DP confirms the complexity of seedling emergence and displays the reason why seedling emergence evaded genetic characterization for so long. The complex quantitative nature of seedling emergence is indicated due to a few major QTLs as well as QTL-by-Environment Interaction (QEI). According to Bernardo (2014), a major marker accounts for >10% of the phenotypic variation. By this definition, a few large effect markers have a range of up to 30% effect on emergence. The identification of a few large effect markers with mainly small effect significant markers by GWAS confirms that seedling emergence is a complex trait controlled by many small-effect loci that may be affected by multiple pleiotropic factors.

### Genome-Wide Association Models

Genetic mapping through association studies has been used to dissect the genetic architecture of various traits in wheat (Adhikari et al. 2012; Lozada et al. 2017, 2019; Bajgain et al. 2019; Lozada and Carter 2020). However, GWAS has not been implemented for deep-sowing seedling emergence in winter wheat but has been conducted in rice (Zhao et al. 2018). Even further, few studies have compared ST-GWAS models, and fewer have compared FarmCPU and BLINK (Liu et al. 2016; Huang et al. 2019). By comparing multiple covariates and GWAS models, we attempted to characterize and dissect a trait with unknown genetic architecture. This allowed us to identify stable markers that are large effect markers with up to 30% effect and 10% R^2^ values. Therefore, we can conclude that these markers are important for seedling emergence and their additivity is noted by an increase in emergence with their accumulation.

The single-locus model used in our study was MLM. The MLM identified significant markers mainly on the 5A chromosome. The MLM has been shown to have difficulties with low power and false positives when identifying significant markers as compared to FarmCPU (Liu et al. 2016). This problem occurs in our study within the DP, and therefore, MLM displays low power in identifying both the major markers and other lower effect significant markers. In our study this was due to high levels of LD between significant markers. The MLM was more consistent in identifying the large effect markers on 5A without the inclusion of the covariates. MLM has been shown to be comparable to FarmCPU for simple traits (Habier et al. 2011; Ward et al. 2019). However, for complex traits such as seedling emergence, the multi-locus models show higher power for small effect markers. The MLM-based models evaluate each genetic marker’s relationship and the overall variation independently; therefore, as the trait increases in complexity and number of pleiotropic effects, the proportion of variation due to a locus decreases (Miao et al. 2019).

FarmCPU and multi-locus models provide the best trade-off between power and false positives (Liu et al. 2016; Miao et al. 2019). The multi-locus models increase the proportion of genetic variance by using the major effect markers as fixed effects and identifying more significant markers (Miao et al. 2019). This is why our multi-locus models were able to dissect the complex trait of seedling emergence accurately and identified both major and minor effect markers. The multi-locus models used in our study were FarmCPU and BLINK and had very similar results in the BL population, but there were fewer significant markers discovered by BLINK than by FarmCPU in the DP. The difference resulted in BLINK not identifying a significant marker in both populations. Other than the markers on 5A that MLM also found, most markers found significant by FarmCPU and BLINK had relatively small effects and R^2^ values indicating multi-locus models’ ability to identify small-effect markers for complex traits.

The majority of markers identified by the MT-GWAS models were also identified in the ST-GWAS models. Further, we had to use the Bonferroni cutoff because the MT-GWAS models displayed inflation of p-values in the QQ plots for the joint analyses. Additionally, the MT-GWAS model uses a single-locus model, which may additionally add to the inflated p-values. Therefore, there was little to no increase in the power for identifying pleiotropic loci compared to the ST-GWAS models. However, the COM model did identify stable markers on chromosome 7A that were not found stable in any ST-GWAS model but were identified in a single year combination. The lack of increase in power for the MT-GWAS models may be due to the lack of phenotypic correlation (Korte et al. 2012). Further, the lack of significant interaction effect loci confirms the positive genetic correlation seen in the majority of years between seedling emergence and coleoptile length. Even though we see a positive genetic correlation, the phenotypic correlation is near zero, indicating a small genetic effect in comparison to the environmental effect. In this scenario, it has been shown that the MT-GWAS models do not outperform ST-GWAS models and confirm why our MT-GWAS models do not outperform the ST-GWAS models (Korte et al. 2012). However, MTMM can still be used as a complement to ST-GWAS models to identify interactions and pleiotropic loci such as the markers on chromosome 7A.

### Covariates

In our study, all of the ST-GWAS and MT-GWAS models used included covariates for population structure as principal components (Price et al. 2006). In addition, the MLM used kinship matrices calculated by VanRaden’s genomic relationship matrix and implemented through GAPIT (VanRaden 2008; Tang et al. 2016). These covariates are commonly used in GWAS studies and allow the models to differentiate genetic and environmental effects (Tibbs Cortes et al. 2021). However, for a complex trait, more confounding associations such as correlated traits or pleiotropic effects may account for a portion of seedling emergence variation (Saltz et al. 2017).

To account for these correlated traits, we compared the use of covariates for Rht markers and coleoptile length. These correlated traits were used due to previous studies reporting large associations between the traits (Allan et al. 1962; Sunderman 1964; Chastain et al. 1995; Schillinger et al. 1998; Botwright et al. 2001; Schillinger 2011). The Rht and coleoptile length covariates had an effect on the multi-locus models but not the MLM. The multi-locus models required the inclusion of covariates in order to identify the major effect of significant markers on chromosome 5A. However, this was not the case for identifying the small effect markers. Therefore, the multi-locus models display a weakness in complex traits to identify both small effect and large effect markers when faced with pleiotropic effects. This signifies that even though modern varieties are no longer dependent on coleoptile length for improved emergence, they still confer pleiotropic effects, confirmed by the results for our MT-GWAS models. We see in the DP that coleoptile length still has a large genotypic correlation depending on the year. Therefore, we recommend using both multi-locus models and covariates for correlated traits to identify both small and large effect loci and increase the power to dissect the genetic architecture of complex traits.

### Association Mapping Populations and Environments

We evaluated the population composition of our populations by using PCs. The DP had a more distinct population structure compared to the BL. This indicates and confirms that the BL population is more genetically related. We used PCs as fixed effects in our GWAS to account for and remove bias due to the presence of underlying population structure in our populations (Price et al. 2006). The closer related the genotypes are, the fewer recombination events, thus preserving marker linkage disequilibrium, and less genetic variation needed to be accounted for by the models (Habier et al. 2007; Bernardo 2020). Due to the increased genetic relatedness and less population structure in the BL, the resolution was increased, and fewer false positives were identified as seen by the large difference in MLM’s significant markers. This may be due to the fact that the BL was purposely selected over generations for deep-sowing seedling emergence in the Washington State University breeding program. The DP on the other hand is composed of varieties from various breeding programs in the Pacific Northwest, with the majority of lines not bred specifically for deep-sowing seedling emergence. Even with this discrepancy, we were still able to identify the stable markers on chromosome 5A in both populations.

The number of environments examined between the two populations differed in addition to the differences between the composition and genetic relatedness of the populations. In the BL, we used a population in a single environment. This restricted our ability to account for GEI in our phenotypic adjustments. However, with the close genetic relationship between the lines, we were still able to identify major effect markers. Whereas in the DP, for both the ST-GWAS and MT-GWAS models, some years had very little significant markers compared to 2015 or the combined years. The year 2015 was important for stable marker identification, with no other individual year identifying the stable markers across populations on chromosome 5A for seedling emergence. Therefore, combining years to partition GEI enabled GWAS models to dissect genetic architectures accurately.

Since seedling emergence is dependent on the environment to create phenotypic variation, it displays GEI. If a trait displays GEI, it follows that so would the QTLs responsible for the phenotypic expression (Bernardo 2020). A change in the ranking of QTLs across environments indicates QEI, and detection of QTLs in some environments, and not others, have been used as a criterion for declaring QEI (Bernardo 2020). Therefore, QEI can be seen with the differing number of significant markers in the different years and the combination of years.

Furthermore, the difficulty in dissecting seedling emergence within and across years can be seen in the discrepancy between the varying genetic and phenotypic correlations between years and traits. The differences in genetic correlations for seedling emergence and coleoptile length from year to year and the near-zero phenotypic correlations can be explained by the large effect of the environment and the multitude of factors that affect seedling emergence. The phenotypic effect can be partitioned into both genetic and non-genetic effects such as the environmental effect (Searle 1961). Since genetic correlation only takes into account the genetic effect, we still see moderate values. The negative value in 2017 can be explained by the different factors that affect seedling emergence, such as coleoptile diameter, force, and speed of emergence. This is further confirmed to be the result of another factor due to the lack of significant markers for the interaction effect in the MT-GWAS models (Korte et al. 2012). As mentioned previously, if the negative genetic correlation was due to the significant loci for coleoptile length, we would see significant markers for the interaction effect (Korte et al. 2012). Further, the near-zero phenotypic correlations display that coleoptile length, while having moderate genetic correlations, has a much smaller genetic effect than the environmental effects such as moisture stress and crusting, and therefore has a little overall phenotypic effect for seedling emergence. In other terms, when genes controlling two traits, such as seedling emergence and coleoptile length, are similar but where the environments pertaining to the expression of the traits have a low correlation. The phenotypic correlation would then be lower than the genetic correlation (Searle 1961).

The inconsistency between years displays the difficulty of dissecting a complex trait affected by multiple factors and heavily dependent on the environment to display phenotypic variation such as seedling emergence. However, the stable markers for seedling emergence on 5A were still identified in both populations across years as well as in both ST-GWAS and MT-GWAS models. Therefore, we can conclude that the genetic architecture’s dissection was successful for both the unselected DP population and selected BL populations.

### Combining favorable alleles

We see a clear increase and additive effect of accumulating favorable alleles in the DP for the stable markers over years. Additionally, there was an additive effect of accumulating favorable alleles for the stable markers over populations in the BL. The difference between the two sets of markers were that the stable markers over populations consisted exclusively of markers on chromosome 5A, whereas the stable markers over years consisted of markers on various chromosomes. This discrepancy shows that the BL may be controlled mainly by the 5A chromosomes with little effect of the other small effect markers on the other chromosomes. Further, seedling emergence for the DP is controlled by both the large-effect and small effect markers on various chromosomes.

The differences in additivity in the populations displays the potential differences between populations purposely selected for complex traits such as seedling emergence and the success of accumulating favorable alleles through traditional phenotypic selection. With the 60 plus years of selection in the BLs, a larger number of favorable alleles may have been accumulated than the DP, consisting of lines from various programs and not centered around deep-sowing emergence selection. Therefore, we can see that through traditional phenotypic selection, the Washington State University breeding program has been able to increase seedling emergence while maintaining the inclusion of Rht alleles and semi-dwarf stature.

Evaluating the cumulative effect of favorable alleles allows us to make assumptions about the types of genetic effects that control seedling emergence. By displaying an increase in seedling emergence with favorable alleles’ accumulation, we can assume that an additive effect conveys seedling emergence. It also shows the ability to pyramid favorable alleles for further improvement of cultivars. Pyramiding favorable alleles has shown to be successful for disease resistance traits (Naruoka et al. 2015; Lewien et al. 2018; Lozada et al. 2019).

### Applications in Breeding

This study compared GWAS models, populations, and covariates in the attempt to dissect the genetic architecture of a complex trait with pleiotropic effects that’s previously been dependent on the environment for phenotypic screening and selection. For these complex traits, including covariates and correlated traits created an advantage in the trait’s dissection. The complex genetic architecture with pleiotropic effects, both small and large effect markers, and the markers’ additivity allowed us to conclude that modern breeding programs no longer are dictated by the large number of studies displaying the dependence of the emergence of dwarf-varieties on coleoptile length. Therefore, this study has also shown empirical results for the ability to breed for correlated traits and overcome perceived genetic disadvantages. In the dryland winter wheat region in Washington, where deep-sowing of winter wheat occurs, plant breeding efforts of 60 years have been able to develop modern varieties that are fast emerging while having the GA insensitive mutant alleles *Rht-B1b* and *Rht-D1b.*

We have also shown the importance of multi-locus models over single-locus models for highly complex traits. The multi-locus models increase in power allows identifying small effect markers located around the genome while accurately identifying the major effect markers as seen in previous comparison (Li et al. 2015; Liu et al. 2016; Miao et al. 2019; Ward et al. 2019; Huang et al. 2019). Further, the MT-GWAS models displayed a lack of power over ST-GWAS models in dissecting correlated traits with low phenotypic correlation as shown in previous studies (Korte et al. 2012).

Even though the large effect markers are advantageous for selecting and improving the trait, identifying smaller effect markers indicates that marker-assisted selection may not be as successful for seedling emergence compared to simpler traits. Therefore, by dissecting the genetic architecture, we can make breeding decisions on the most accurate method for rapid improvements, such as the inclusion of genomic selection or phenotypic selection that may be more accurate for substantial genetic gain (Bernardo 2008). For instance, our GWAS models conclude there are only a few large effect markers while identifying mainly small effect markers. The use of genomic selection may be more successful while still having the capability of including large effect markers as fixed effects in the genomic selection models. As in this case, the markers convey effects of 30% for seedling emergence and display additive effects. These high effects meet the criteria for inclusion set forward by Bernardo (2014). Even when GWAS and dissections may not provide results in identifying candidate genes or large effect markers for use in selection, it can allow us to conclude the most effective method of selection, which allows breeders to be more efficient with their limited resources while meeting their program’s goals and targeted genetic gains.

## Conclusion

This study displayed the ability for GWAS to dissect the genetic architecture of a complex trait such as deep-sown seedling emergence. Both ST-GWAS and MT-GWAS models identified a few large effect and many small effect markers for seedling emergence. Additionally, neither the ST-GWAS or MT-GWAS models identified large pleiotropic effect markers between seedling emergence and coleoptile length. The ST-GWAS and MT-GWAS models did not identify the same significant markers for seedling emergence and coleoptile length, and the MT-GWAS models did not identify any interaction effect markers. However, by using multi-locus models in conjunction with covariates for correlated traits, we were able to increase power over single-locus models to dissect a complex traits in both the DP and BL populations. Additionally, the DP displayed the necessity for combining years for consistent identification of significant markers for a trait dependent on the environment for phenotypic variation. Further, the MT-GWAS models displayed a lower power over ST-GWAS models for single-trait analysis and inflated p-values for joint analysis but still identified the large effect markers on 5A. Further, the MT-GWAS model uses a single-locus model, which may add to the inflated p-values. Therefore, the MT-GWAS models displayed a lack of power in dissecting correlated traits with low phenotypic correlation. Therefore, by using multi-locus models combined with pleiotropic covariates, breeding programs can uncover the complex nature of traits to help identify candidate genes and the underlying architecture of a trait to make more efficient breeding decisions and selection methods.

## Declarations

### Funding

This research was partially funded by the National Institute of Food and Agriculture (NIFA) of the U.S. Department of Agriculture (Award number 2016-68004-24770), Hatch project 1014919, and the O.A. Vogel Research Foundation at Washington State University.:

### Conflicts of Interest

The authors declare that the research was conducted in the absence of any commercial or financial relationships that could be construed as a potential conflict of interest.

### Availability of data and material

The datasets generated for this study can be found at https://github.com/lfmerrick21/GWAS-Complex-Traits

### Author’s Contributions

LM: conceptualized the idea, analyzed data, and drafted the manuscript; AB: genotyped the KASP markers, reviewed, and edited the manuscript; ZZ: reviewed and edited the manuscript; AC: supervised the study, conducted field trials, edited the manuscript, and obtained the funding for the project.

## Acknowledgments

The authors would like to acknowledge the Washington State University Winter Wheat Breeding Program personnel Gary Shelton and Kyall Hagemeyer for plot maintenance and data collection under field conditions and Kerry Barlow for coleoptile length data collection. We would also like to thank Gina Brown-Guedira, Jared Smith, Brian Ward and staff at the Eastern Regional Small Grains Genotyping Laboratory for their assistance with DNA library prep and GBS sequencing and analysis.

## Notes

https://github.com/lfmerrick21/GWAS-Complex-Traits

### Competing Interest Statement

The authors have declared no competing interest.

### Summary of Updates

Fixed Table and Figure Captions and Numbering.

