## Supplementary Material for "Comparison of Single-Trait and Multi-Trait Genome-Wide Association Models and Inclusion of Correlated Traits in the Dissection of the Genetic Architecture of a Complex Trait in a Breeding Program"

| **Table S1** Coleoptile length Cullis heritability and trial statistics for the diversity panel (DP) population phenotyped from 2014 to 2016 | | | | | | |
| --- | --- | --- | --- | --- | --- | --- |
| Population | Heritability | Max (mm)^a^ | Mean (mm) | Min (mm)^b^ | SD^c^ | CV^d^ |
| DP | 0.89 | 123 | 82 | 67 | 7 | 8.70% |
| ^a^Max: maximum  ^b^Min: minimum  ^c^SD: standard deviation  ^d^CV: coefficient of variation | | | | | | |

| **Table S2** Analysis of variance (ANOVA) and gentoype-by-environment analysis for deep-sowing seedling emergence in individual and combined trials for the diversity panel (DP) population and breeding line (BL) population phenotyped from 2015 to 2019 and 2015, respectively | | | | | | | | |
| --- | --- | --- | --- | --- | --- | --- | --- | --- |
| Population | Set | Year | Check (Fixed/  Estimate) | Genotype (Random/  Variance) | Block (Random/  Variance) | Environment (Random/  Variance) | GEI  (Random/  Variance) | Residual |
| DP | DP | 2015 | 17.21 | 161.90 * | 171.50 *** |  |  | 161.30 |
| DP | DP | 2017 | 3.56 | 45.28 * | 7.07 *** |  |  | 67.22 |
| DP | DP | 2018 | -0.05 | 149.76 | 0.00 |  |  | 50.00 |
| DP | DP | 2019 | -2.76 | 190.70 | 343.10 *** |  |  | 325.80 |
| DP | DP | 2015-2017 | 17.22 | 31.76 ** | 22.95 | 14.37 | 69.93 | 116.04 |
| DP | DP | 2015-2018 | 17.27 | 17.74 ** | 26.40 | 197.91 | 79.41 * | 114.77 |
| DP | DP | 2015-2019 | 17.34 | 25.36 *** | 25.05 | 43.27 | 118.83 * | 144.24 |
| BL | F_3:5_ | 2015 | 11.22 | 164.80 *** | 109.40 *** |  |  | 142.50 |
| 0 ‘***’ 0.001 ‘**’ 0.01 ‘*’; DH: doubled-haploid; GEI: genotype-by-environment interaction | | | | | | | | |

| **Table S3** Phenotypic correlations for deep-sowing seedling emergence adjusted means between individual and combined trials for the diversity panel (DP) population and breeding line (BL) population phenotyped from 2015 to 2019 and 2015, respectively | | | | | | | |
| --- | --- | --- | --- | --- | --- | --- | --- |
|  | DP 2017 | DP 2018 | DP 2019 | DP 2015-2017 | DP 2015-2018 | DP 2015-2019 | BL 2015 |
| DP 2015 | 0.50*** | 1.00 | 0.19* | 0.87*** | 0.94*** | 0.65*** | 0.93 |
| DP 2017 |  | 0.50 | 0.94 | 0.87*** | 0.76*** | 0.98*** | 0.78* |
| DP 2018 |  |  | 0.19* | 0.87 | 0.94*** | 0.65*** | 0.93 |
| DP 2019 |  |  |  | 0.65* | 0.50*** | 0.87*** | 0.54 |
| DP 2015-2017 |  |  |  |  | 0.98*** | 0.94*** | 0.99 |
| DP 2015-2018 |  |  |  |  |  | 0.87*** | 1.00 |
| DP 2015-2019 |  |  |  |  |  |  | 0.89 |
| 0 ‘***’ 0.001 ‘**’ 0.01 ‘*’ | | | | | | | |

| **Table S4** The frequency of the Rht alleles in the diversity panel (DP) population and breeding line (BL) population for *Rht-B1b* and *Rht-D1b* | | | |
| --- | --- | --- | --- |
| DP | Rht | Number of Lines | Frequency |
|  | *Rht-B1b* | 267 | 0.564 |
|  | *Rht-D1b* | 164 | 0.347 |
|  | *Rht-B1b* Heterozygous | 2 | 0.004 |
|  | *Rht-D1b* Heterozygous | 5 | 0.011 |
|  | *Rht-D1b* with a Heterozygous *Rht-B1b* | 22 | 0.047 |
|  | *Rht-B1b* with a Heterozygous *Rht-D1b* | 0 | 0 |
|  | *Rht-B1b* and *Rht-D1b* Heterozygous | 13 | 0.027 |

| **Table S5** Frequencies of the favorable alleles for the stable markers identified across populations and years in the diversity panel (DP) population and breeding line (BL) population | | | | |
| --- | --- | --- | --- | --- |
| Population | Significance | Marker | n | frequency |
| DP | Years | S2A_605310547 | 305 | 0.64 |
|  | Years | S2A_748461724 | 446 | 0.94 |
|  | Years | S2B_301162652 | 248 | 0.52 |
|  | Years | S2B_403420819 | 471 | 1.00 |
|  | Years | S2B_789673174 | 190 | 0.40 |
|  | Pops and Years | S5A_514260464 | 432 | 0.91 |
|  | Pops and Years | S5A_522153944 | 456 | 0.96 |
|  | Pops and Years | S5A_523025549 | 454 | 0.96 |
|  | Years | S5A_523147137 | 464 | 0.98 |
|  | Years | S5A_535147597 | 323 | 0.68 |
|  | Years | S5B_491273019 | 386 | 0.82 |
|  | Years | S7A_82946533 | 107 | 0.23 |
|  | Years | S7B_663828309 | 203 | 0.43 |
| BL | Years | S2A_605310547 | 171 | 0.62 |
|  | Years | S2A_748461724 | 242 | 0.88 |
|  | Years | S2B_301162652 | 202 | 0.73 |
|  | Years | S2B_403420819 | 276 | 1.00 |
|  | Years | S2B_789673174 | 55 | 0.20 |
|  | Pops and Years | S5A_514260464 | 223 | 0.81 |
|  | Pops and Years | S5A_522153944 | 266 | 0.96 |
|  | Pops and Years | S5A_523025549 | 267 | 0.97 |
|  | Years | S5A_523147137 | 272 | 0.99 |
|  | Years | S5A_535147597 | 181 | 0.66 |
|  | Years | S5B_491273019 | 197 | 0.71 |
|  | Years | S7A_82946533 | 38 | 0.14 |
|  | Years | S7B_663828309 | 130 | 0.47 |
| n: Number of lines | | | | |

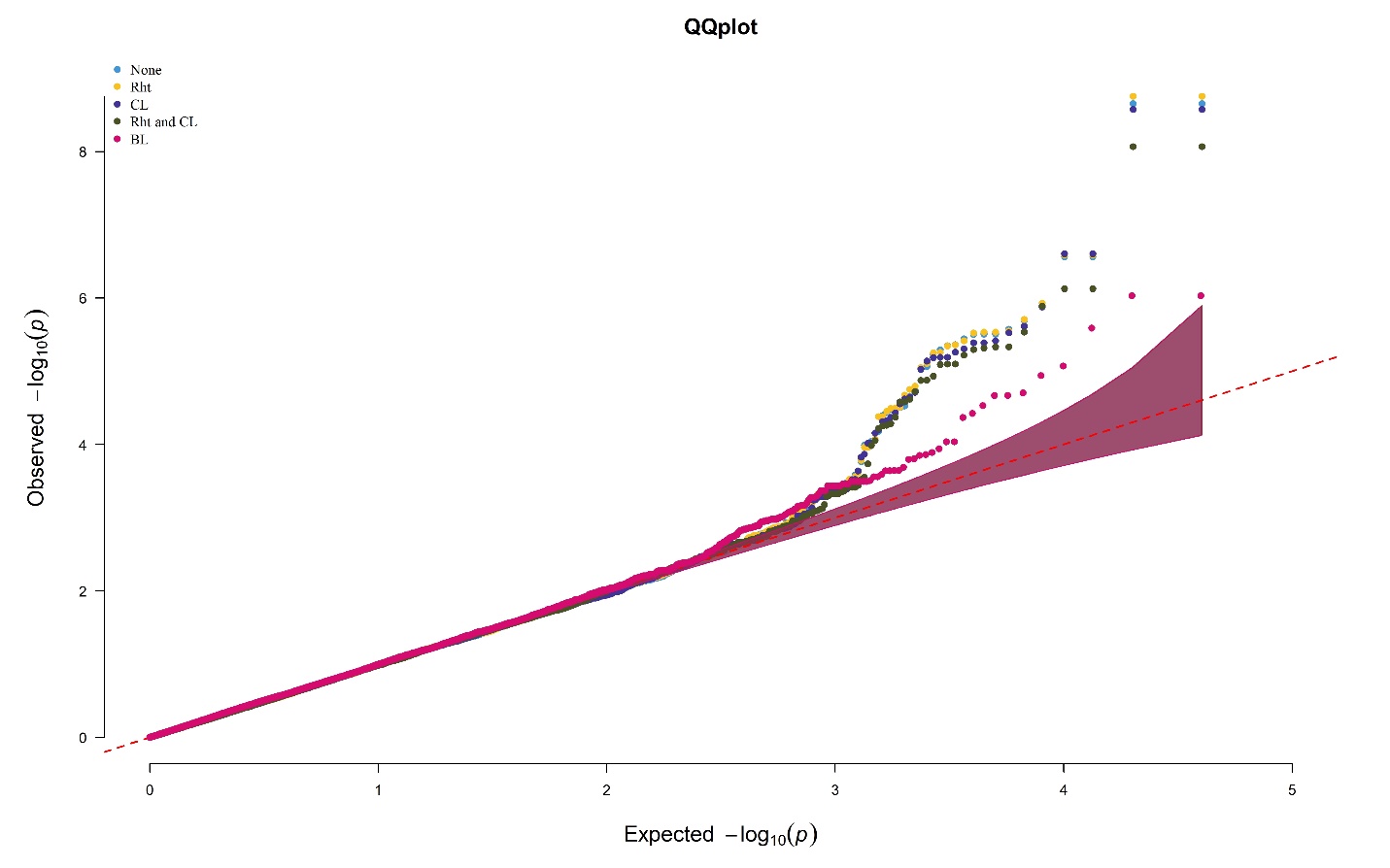

**Fig. S1** Quantile-Quantile (QQ) plots comparing the genome-wide association mixed linear model with the inclusion of no covariates, Rht alleles, Coleoptile Length, Rht alleles and Coleoptile Length within the diversity panel and without covariates in the breeding line trial. The models were conducted for deep-sowing seedling emergence in a Pacific Northwest winter wheat diversity panel phenotyped across trials from 2015-2018 and a breeding line trial phenotyped in 2015 in Lind, WA

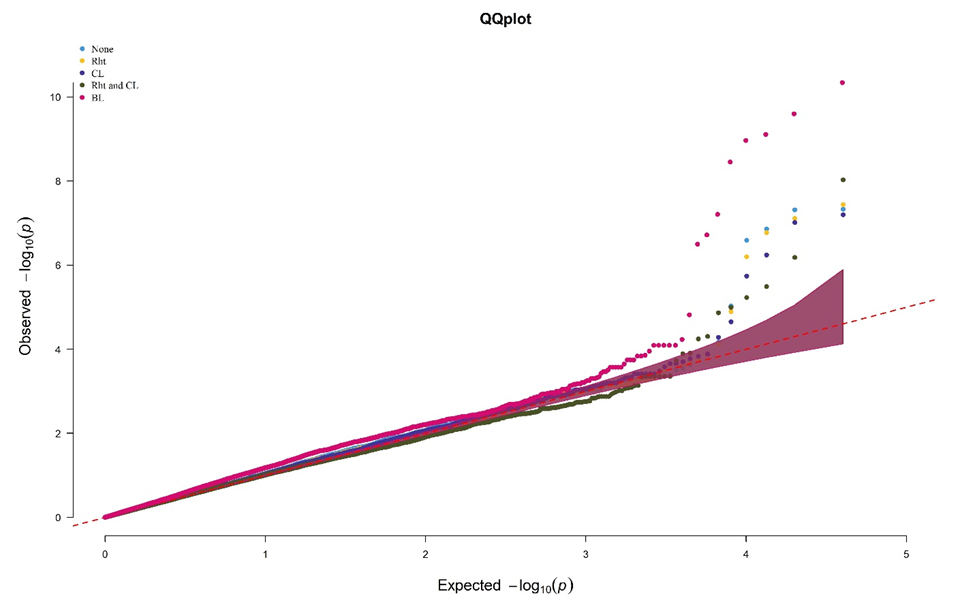

**Fig. S2** Quantile-Quantile (QQ) plots comparing the genome-wide association model, BLINK, with the inclusion of no covariates, Rht alleles, Coleoptile Length, Rht alleles and Coleoptile Length. The models were conducted for deep-sowing seedling emergence in a Pacific Northwest winter wheat diversity panel phenotyped across trials from 2015-2018 and a breeding line trial phenotyped in 2015 in Lind, WA

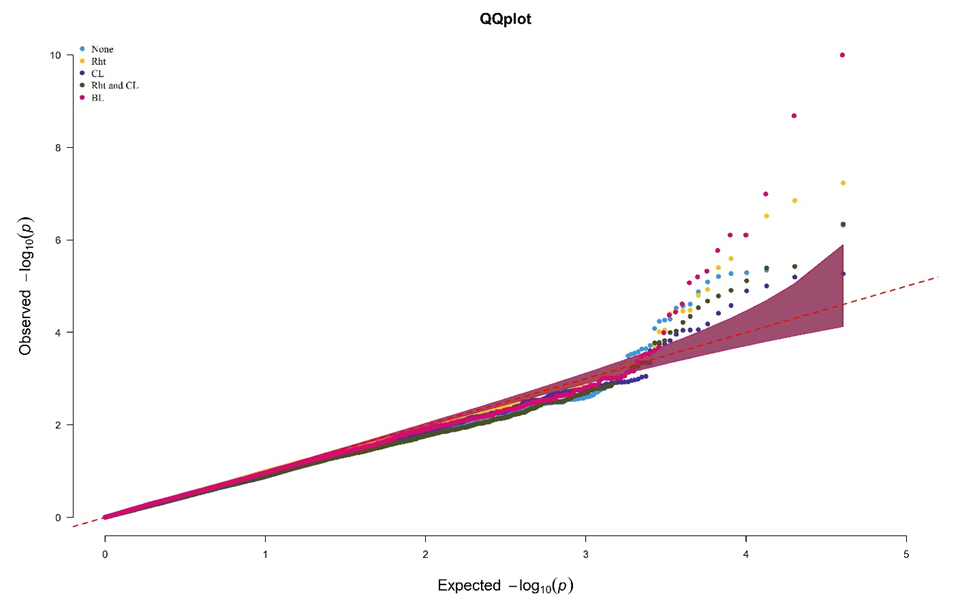

**Fig. S3** Quantile-Quantile (QQ) plots comparing the genome-wide association model, FarmCPU, with the inclusion of no covariates, Rht alleles, Coleoptile Length, Rht alleles and Coleoptile Length. The models were conducted for deep-sowing seedling emergence in a Pacific Northwest winter wheat diversity panel phenotyped across trials from 2015-2018 and a breeding line trial phenotyped in 2015 in Lind, WA

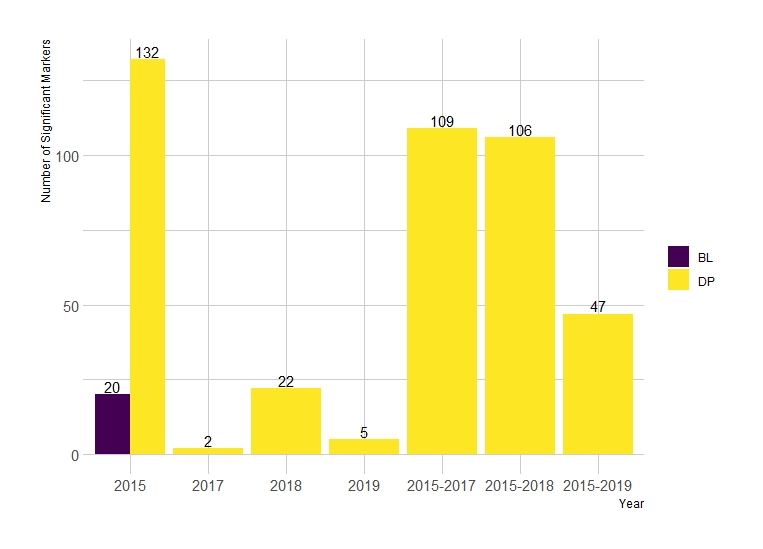

**Fig. S4** The sum of significant markers identified in individual years and combination of years across both the diversity panel (DP) and breeding line (BL) populations from all combinations of single-trait models

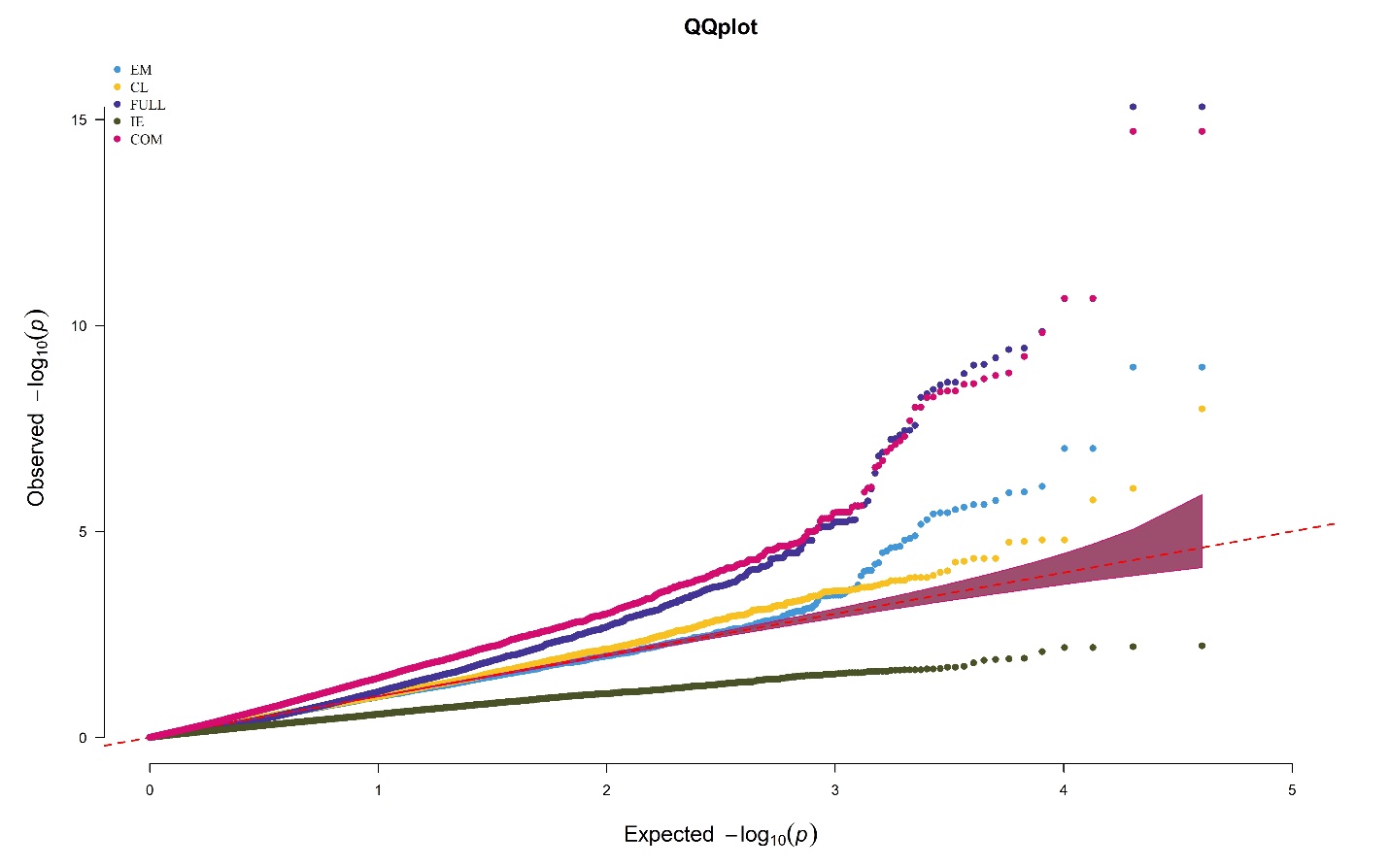

**Fig. S5** Quantile-Quantile (QQ) plots comparing the multi-trait genome-wide association model (MT-GWAS), for seedling emergence (EM), coleoptile length (CL), full model (FULL), interaction effect model (IE), and common model (COM). The models were conducted on a Pacific Northwest winter wheat diversity panel phenotyped across trials from 2015-2018 in Lind, WA

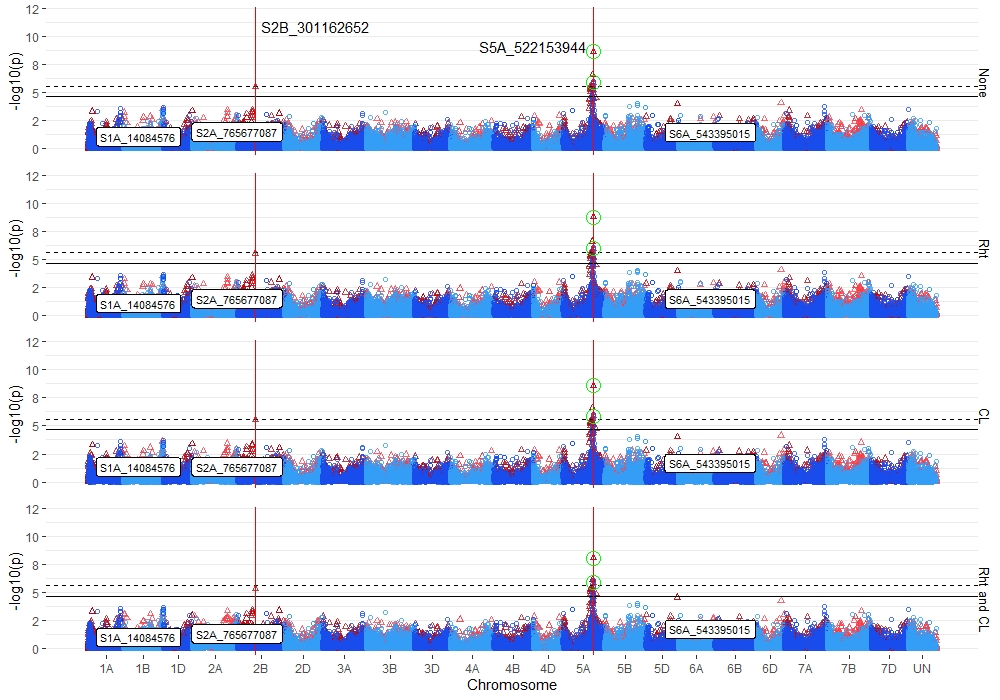

**Fig. S6** Stacked manhattan plots for genome-wide association studies (GWAS) using MLM for identifying significant loci controlling deep-sowing seedling emergence in a Pacific Northwest winter wheat diversity panel phenotyped across trials from 2015-2018 and a breeding line (BL) trial phenotyped in 2015 in Lind, WA. Covariates used were Reduced Height markers (Rht) and BLUPs for coleoptile length (CL). Red triangles display GWAS results in the DP, and blue circles identify GWAS results in the BL. Significant markers using an FDR cutoff with an alpha=0.05 are placed above the solid black line for the DP and the dashed black line for the BL. Significant markers across three models are highlighted with a solid red vertical line, and significant markers across two models are highlighted with a dashed green vertical line identifying their positions. Significant markers across both populations are encircled in a green circle. Markers enclosed in a white text box display significant markers identified in GWAS studies for coleoptile length

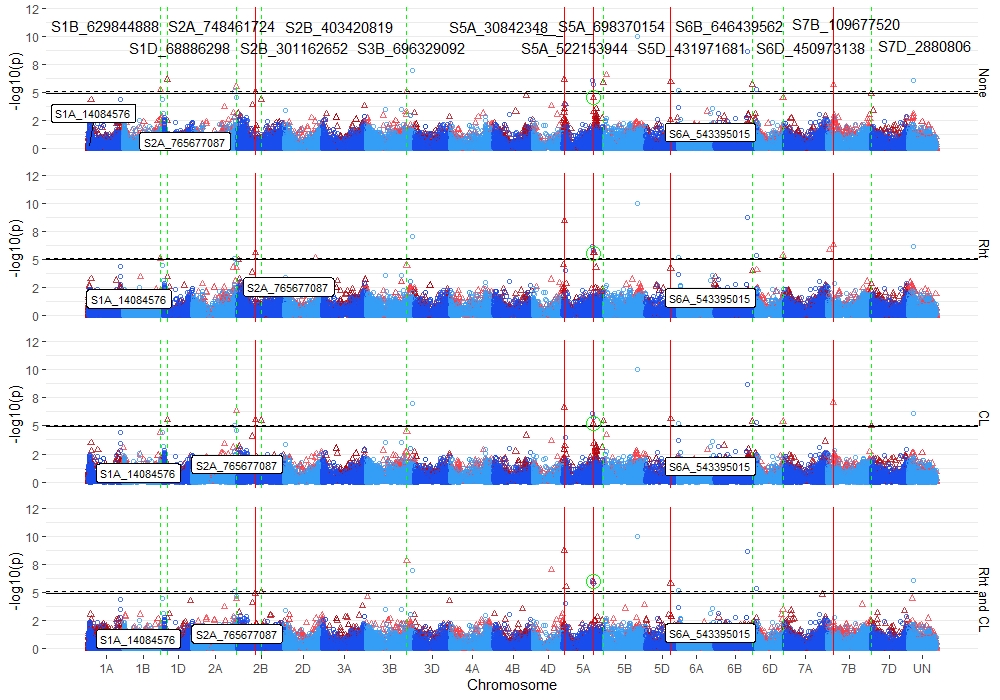

**Fig. S7** Stacked manhattan plots for genome-wide association studies (GWAS) using FarmCPU for identifying significant loci controlling deep-sowing seedling emergence in a Pacific Northwest winter wheat diversity panel phenotyped across trials from 2015-2019 and a breeding line (BL) trial phenotyped in 2015 in Lind, WA. Covariates used were Reduced Height markers (Rht) and BLUPs for coleoptile length (CL). Red triangles display GWAS results in the DP, and blue circles identify GWAS results in the BL. Significant markers using an FDR cutoff with an alpha=0.05 are placed above the solid black line for the DP and the dashed black line for the BL. Significant markers across three models are highlighted with a solid red vertical line, and significant markers across two models are highlighted with a dashed green vertical line identifying their positions. Significant markers across both populations are encircled in a green circle. Markers enclosed in a white text box display significant markers identified in GWAS studies for coleoptile length

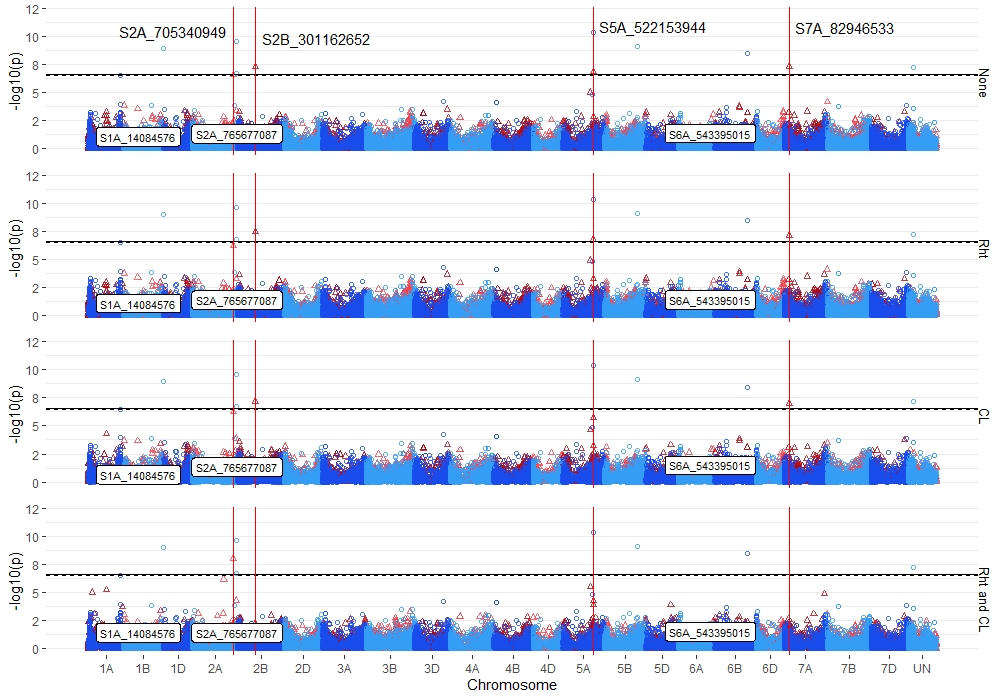

**Fig. S8** Stacked manhattan plots for genome-wide association studies (GWAS) using BLINK for identifying significant loci controlling deep-sowing seedling emergence in a Pacific Northwest winter wheat diversity panel phenotyped across trials from 2015-2018 and a breeding line (BL) trial phenotyped in 2015 in Lind, WA. Covariates used were Reduced Height markers (Rht) and BLUPs for coleoptile length (CL). Red triangles display GWAS results in the DP, and blue circles identify GWAS results in the BL. Significant markers using an FDR cutoff with an alpha=0.05 are placed above the solid black line for the DP and the dashed black line for the BL. Significant markers across three models are highlighted with a solid red vertical line, and significant markers across two models are highlighted with a dashed green vertical line identifying their positions. Significant markers across both populations are encircled in a green circle. Markers enclosed in a white text box display significant markers identified in GWAS studies for coleoptile length
